# Difference in the impacts of the neonicotinoid dinotefuran administered through sugar syrup from that through pollen paste on a honeybee colony in the long-term field experiment

**DOI:** 10.1101/014779

**Authors:** Toshiro Yamada, Yasuhiro Yamada, Kazuko Yamada

## Abstract

**Summary:** We have previously examined the impact of neonicotinoid pesticides, dinotefuran and clothianidin, on honeybee Apis mellifera colonies in long-term field experiments when they were simultaneously administered through both vehicles of sugar syrup and pollen paste (Yamada *et al.*, 2012). The independent effect of a pesticide through two vehicles has not been studied in our previous work. In this paper, we investigated the independent impact of dinotefuran through each of the two vehicles. We confirmed that dinotefuran intake per bee until colony extinction due to administration through pollen paste (DF-TIPP) was roughly one-fifth as much as that through sugar syrup (DF-TISS). The intake was largely independent of dinotefuran concentration. We considered the possibility of DF-TIPP per bee as an indicator to assess the impact of persistent pesticide on a honeybee colony in a practical apiary.

This work has replicated the finding that a honeybee colony has dwindled away to nothing after assuming an aspect of a colony collapse disorder (CCD) by administration of the neonicotinoids dinotefuran and clothianidin in our previous work, regardless of the vehicles. In addition, a failure in wintering was observed in case of administration of dinotefuran with the lowest concentration in this work even if the colony appeared vigorous before winter.

We can infer that a CCD and a failure in wintering may have the same roots of chronic toxicity of neonicotinoids under conditions of low concentrations due to the persistency and high toxicity which are characteristic of them.

## Introduction

A rapid decline in honeybee colonies which has been first reported in Europe is becoming a serious worldwide social problem. (Cox-Foster *et al.*, 2007; Potts *et al.*, 2010; Van Engelsdorp el al., 2012; Lebuhn *et al.*, 2013). A trend analysis of US agriculture demonstrated that a diminution in managed or wild pollinator populations could seriously threaten crop production (Calderone, 2012). Massive losses of honeybee colonies have begun to be actively investigated in the world when mysterious phenomenon inexplicable by physiological changes of a honeybee colony was called a colony collapse disorder (CCD) which is characterized by the rapid loss of adult bee populations from a honeybee colony with little dead bees near the hive and the state at the final stages of collapse with the presence of capped brood, a queen, a few adult bees and food stores (both honey and bee pollen). And massive losses of honeybee colonies by a failure in wintering also have threatened beekeepers and agricultural workers (Steinhauer *et al.*, 2014; van der Zee *et al.*, 2014).

Various factors such as pesticides, mites, viruses, stresses, poor nutrition and weather patterns have been postulated as causes on a CCD and a failure in wintering (van der Sluijs *et al.*, 2013). Neumann and Carreck (2010) have reviewed fact-finding results on honeybee colony losses in the world and their causal factors. In USA, pesticides applied to crops and those, with their residues, used in apiculture in hive products were reviewed and examined to determine what could play a role in a CCD and other colony problems (Johnson *et al.*, 2010).

Recently, it have become predominantly convincing that a CCD and a failure in wintering are caused by a systemic pesticide itself such as a neonicotinoid and a synergy between the pesticide and others such as mites (van der Sluijs *et al.*, 2013).

Many studies have been reported on the impact of neonicotinoid pesticides on a honeybee colony under laboratory conditions or controlled field-experimental conditions as follows: Sublethal oral doses of imidacloprid decreased the fecundity of worker bumblebees of queenless microcolonies, showing a dose dependence that principally correlated with nutrient limitations imposed by antifeedant effects (Laycock *et al.*, 2012). Investigation of honeybee foraging behavior by the radiofrequency identification method showed that sublethal oral administration of clothianidin and imidacloprid impacted the flight frequency and duration of flight activity (Schneider *et al.*, 2012). Honeybee behavior influenced by a sublethal oral dose of imidacroprid and acaricide were effectively measured by the video-tracking method (Teeters *et al.*, 2012). Although a sublethal dose of imidacloprid had no effect on capped brood, pupation, and eclosion rates of honeybee larvae, the proboscis extension reflex test after emergence revealed impairment of the development of olfactory ability (Yang *et al.*, 2012). Assessment of the effects of imidacloprid ingestion by stingless bee larvae on their survival, development, neuromorphology, and adult walking behavior revealed that these larvae were particularly susceptible to imidacloprid because the pesticide caused both high mortality and sublethal effects that impaired brain development and compromised mobility at the young adult stage (van Tomé *et al.*, 2012).

Pseudo-field testing revealed that the foraging activity of honeybees decreases with pesticide concentrations of a few micrograms/kilograms (imidacloprid, fipronil) (Colin *et al.*, 2004), and sublethal doses of imidacloprid are shown to affect the foraging behavior of honeybees (Yang *et al.*, 2008). Furthermore, a particular vulnerability of honeybee behavior to sublethal doses of acetamiprid was suggested under laboratory conditions (El Hassani *et al.*, 2008). Under field conditions, a difference was observed in the survival rate between a group of bees treated with 70 ng imidacloprid and a control group (Visser and Blacquière, 2010). The sudden death phenomenon of bees during sowing suggests the synergistic effect of high humidity and toxicity of a powder containing neonicotinoids (Marzaro *et al.*, 2011). Colonies of bumblebees treated with field-realistic levels of imidacloprid showed a significantly reduced growth rate and a production rate of new queens about 15% that of the control colonies (Whitehorn *et al.*, 2012). Sublethal exposure of honeybees to thiamethoxam at levels that could put a colony at risk of collapse caused high mortality due to homing failure (Henry *et al.*, 2012), and Matsumoto (2013) demonstrated that neonicotinoid and pyrethroid exposure reduced successful homing flights at doses far below the median lethal dose (LD_50_) in the field where neonicotinoid caused the reduction at relatively lower exposure than pyrethroid. Chronic exposure of bumblebees to imidacloprid at approximate field-level concentrations reduced the amount of pollen collected because of impaired foraging efficiency and (Gill *et al.*, 2012).

Only a few studies have been reported on the impact of neonicotinoid pesticides on a honeybee colony under long-term field conditions (Lu *et al.*, 2012; 2014: Yamada *et al.* 2012). Yamada *et al.* (2012) have tried to verify the possibility of neonicotinoids, clothianidin and dinotefran, causing a CCD based on experiments in 2010 in which a CCD was reproduced by field testing. The experiments were carried out at three concentrations (low, middle, high) for each of clothianidin and dinotefuran and similar results were obtained for clothianidin and dinotefuran. At low and middle concentrations each pesticide was continuously administered through both vehicles (foods) of sugar syrup and pollen paste till the colony extinction. At high concentration it was not administered till the colony extinction after it was administered through both vehicles only for the first time around. At high concentration massive dead bees were found near the hive of each high concentration colony just after the first administration of each pesticide; thereafter, a few dead bees were found similarly to control colonies and experimental colonies at low and high concentration of pesticide. A queen existed in the hive together with some adult bees just before the cony extinction. Each number of adults, capped brood and dead bees in the colony where dinotefuran was administered similarly changed with day to that of clothianidin while assuming an aspect of a CCD. In addition, they have made it clear from the speculations with the NMR spectral analysis that dinotefuran is thermally and radiationally stable under the conditions of 50 °C×24hrs or 310nm×50 W/m^2^. It has been suggested that neonicotinoids are very persistent and continue to accumulate in organisms. Around the same time frame as our previous paper, Lu *et al.* (2012) have reported that the neonicotinoid imidacloprid can cause a failure in wintering on honeybee colonies through field experiments in several apiaries and have recently confirmed their previous results by a long-term field experiments with imidacloprid and clothianidin (Lu *et al.*, 2014).

Yamada *et al.* (2012) have inferred that the bottom cause of the inexplicable collapse of honeybee colonies is probably neonicotinoid pesticides which are persistent, systemic and highly toxic for the reasons why the characteristics of a neonicotinoid pesticide lead to the disorientation of honeybees, the reduction in ovipositional performance of a queen and the infestation of mites and viruses due to the weakening of honeybees when exposed to honeybees in its low concentrations for a long period of time, or their instant death when exposed to them in its high concentrations. One of convincing causal factors is probably neonicotinoid pesticides themselves or factors compounded of neonicotinoid pesticides and others such as mites (Alaux *et al.*, 2010; Pettis *et al.*, 2012). That is, neonocotinoid pesticides seem to play a vital part in massive losses of honeybee colonies. An extensive survey of honeybee colony losses was conducted in Japan in 2008-2010, which revealed that bee loss was mostly due to neonicotinoids sprayed for stink bug control following rice flowering (Taniguchi *et al.*, 2012).

In previous studies a pesticide was administered to a honeybee colony through sugar syrup or its analogues containing a given amount of pesticide (Colin *et al.*, 2004; Laurino *et al.*, 2011; Henry *et al.*, 2012; Schneider *et al.*, 2012; Lu *et al.*, 2012, 2014; Teeters *et al.*, 2012), or was simultaneously administered through both sugar syrup and pollen paste (Yamada *et al.*, 2012). The previous studies have paid never attention to the intake route of a pesticide by honeybees though there can be a difference in the influence of a pesticide on a honeybee colony between different intake routes of sugar syrup and pollen paste.

It is known that brood and a queen preferentially take pollen which is an important protein source for honeybees rather than honey which is an important energy source for them though all members of a honeybee colony take both. Therefore, toxic honey with neonicotinoid pesticides may affect a honeybee colony differently from toxic pollen. We cannot find papers on the influence of the route, through which honeybees takes a pesticide, on a honeybee colony.

In this work we therefore clarify the influence of dinotefuran, which is the most commonly used neonicotinoid in Japan, on honeybee colonies depending on a kind of the administration vehicle (sugar syrup or pollen paste) under the field experiments performed from July 9^th^ in 2011 to April 2^nd^ in 2012. Experimental concentrations of dinotefuran were determined with reference to clothianidin content of 5 ppm in water detected near a paddy field where the pesticide was crop-dusted (Kakuta *et al.*, 2011), similarly to the previous work (Yamada *et al.*, 2012). The neonicotinoid dinotefuran was administered through either sugar syrup or pollen paste in a hive at two concentrations each, on the assumption that sugar syrup corresponds to nectar or honey and water and pollen paste does to bee bread in the natural environment. By the way, in the natural environment we must pay attention to the possibility that we cannot always make a clear-cut distinction between the influence of a pesticide through honey (nectar) and that through bee bread because bee bread is made by mixing pollen with water and nectar from the bee’s mouth, which causes the pollen granules to grow. Even if pollen is not contaminated by a pesticide, the bee bread may be contaminated by it through toxic nectar or water in the natural environment. We can use the results in this work with due consideration for the above possibility in the natural environment.

In addition to the comparison between two intake routes of the pesticide, we confirm in this study the fact that a honeybee colony have become extinct to nothing after assuming an aspect of a CCD in the previous experiments (Yamada *et al.*, 2012) and we discuss the possibility for an indicator to assess a collapse of a honeybee colony due to persistent pesticides such as neonicotinoids instead of LD_50_ which is related to the death of an individual honeybee.

## Materials and Methods

### Ethics statement

We clearly state that no specific permissions were required for these locations/activities because the apiary at which we performed the experiments for this study belongs to the author (Toshiro Yamada). We confirm that the field studies did not involve endangered or protected species.

### Materials and preparation of pesticide concentrations

The pathways where honeybees take pesticides from nectar, pollen, water and so on into a colony are very complicated. For examples, nectar and pollen contaminated with pesticides are imported from fields into a hive, most of which are stored on combs as honey or bee bread after the pesticides are diluted with pesticide-free one and have an enduring effect on a honeybee-colony. Toxic water is fed to young bees and brood in the early spring or is used to reduce the temperature of cells in summer and result in contamination of the whole hive with pesticides. Field experiments include many uncontrollable factors contrary to laboratory ones. In order to decrease in ambiguity as much as possible we try to .avoid unintentional contamination with pesticides. First we selected an experimental apiary site where there were no large paddy fields and orchards in the vicinity where aerial-spraying supposed to be conducted, whose pesticide-concentration is about 100 times as high as hand-spraying, (for example, the concentration of dinotefuran is 12500 ppm in aerial-spray solution in Japan and 1000 ppm in hand-spray one for extermination of stinkbugs). Secondly, we located a honeybee-watering place in the experimental apiary where leaf mustard *Brassica juncea* and hairy vetch *Vicia villosa* were planted to supply experimental honeybee colonies with pesticide-free water nectar and pollen and to minimize the effects of environmental factors.

Experiments were performed in 2011 under experimental conditions as tabulated in Table 1. STARKLE MATE^®^ (10% dinotefuran; Mitsui Chemicals Aglo, Inc., Tokyo, Japan) which is a commercial product and mostly sprayed on rice paddies in Japan was used in this study instead of dinotefuran only in order to bring the experimental conditions closer to the realistic field ones. STARKLE MATE^®^ includes auxiliary materials such as stabilizers, surfactants and adjuvants which are assumed to be biologically inert. Though the auxiliary materials may slightly affect honeybee colonies (Ciarlo *et al.*, 2012), we have expressed the experimental concentration of the active ingredient of the pesticide by the concentration of dinotefuran only in this study.

**Table 1.** Details of experimental methods in this work

|  | **Details of Experimental methods** |
| --- | --- |
| **Experimental period** | July 9 in 2011 to April 2 in 2012 |
| **Circumstnces in an apiary** | Without the large paddy field within a radius of two kilometers of the apiary and controllable time and place for crop-dusting |
| **Confirmation of experiment** | Double-checked by two persons |
| **Number of hives** | Five hives (one control & four dose tests) |
| **Initial composition of a hive** | three combs & a feeder |
| **Initial number of honeybees** | ca. 3,000 |
| **Record of colony conditions** | Photos of all combs and the inside of a hive with honeybees and all combs without honeybees taken in every experiment |
| **Record of entrance circumstance** | Time-lapse photography at the interval of 1 hour with two cameras |
| **Number of adult bees in a hive** | Directly counted with a counter on photos of all combs and the inside of a hive one by one |
| **Number of brood in a hive** | Directly counted with a counter on photos of all combs without honeybees |
| **Number of dead bees** | Directly counted one by one with distinguishing between dead bees in a hive and those on the tray installed under a hive |
| **Intake of pesticide of honeybees** | Accurately weighed by a weighing instrument |
| **A kind of pesticide** | Dinotefuran (DF) which is the top amount of pesticide used in Japan |
| **Concentration of pesticide administered into a colony via sugar syrup or pollen paste** | 1 (DF-Low) and 10 ppm (DF-High) of DF in sugar syrup. As pollen paste is made of sugar syrup and pure pollen in a ratio of 1.3:1 after pulverizing pollen into finer patticles, the concentration of pesticide in pollen paste becomes 0.565 ppm for DF-Low/Pollen and 5.65 ppm for DF-High/Pollen. Sugar syrup was made of an equal amount of sugar and water. |
| **Administration method of pesticide** | 1) Separate administration of either sugar syrup or pollen paste with pesticide in each hive<br>2) For DF-High, the pesticide was administered only in the beginning of experiment (that is, the period of administration was seven days) and for the others, it continued to be administered during the experimental period till the colony become extinct |
| **Prevention of swarming** | Experiment start after the swarming period and then addition of a new comb and change to a bigger hive |
| **Confirmation of a queen bee** | Record by photos |
| **Flowering site** | Flowering site was installed in the apiary such as the fields of leaf mustard Brassica juncea and hairy vetch Vicia villosa. |
| **Honeybee watering place** | New watering place was provided for honeybees in the apiary |
| **Hornet catcher** | Installation of a hornet catcher in each hive after summer |
| **Starting time of each experiment** | Early morning if possible |
| **Others** | Record by photos about troubles such as wax worms, bee-beetles, etc |

Based on a concentration of 5 ppm clothianidin detected near a paddy field in which the pesticide was crop-dusted (Kakuta *et al.*, 2011) and maximum residue limits (MRLs) of agricultural chemicals in foods in Japan (JFCRF, 2013) where MRLs of dinotefuran range from 0.1 ppm to 25 ppm, those of clothianidin from 0.02 ppm to 50 ppm and those of imidacloprid from 0.05 ppm to 10 ppm, the highest concentration of dinotefuran administered to a colony was determined to be 10 ppm, because the insecticidal activity of dinotefuran for stink bugs is about 0.4 times that of clothianidin from our previous work (Yamada *et al.*, 2012) and five ppm of clothianidin is equivalent to 12.5 ppm of dinotefuran. The concentrations of dinotefuran used were 1 and 10 ppm (termed Low and High, respectively) in sugar syrup and 0.565 and 5.65 ppm (termed Low and High, respectively) in pollen paste, because pollen paste comprises pure pollen without pesticide and sugar syrup with dinotefuran in the ratio 1:1.3. We can now define DF-Low/Syrup, DF-Low/Pollen, DF-High/Syrup, and DF-High/Pollen as sugar syrup with 1 ppm dinotefuran, pollen paste with 0.565 ppm dinotefuran, sugar syrup with 10 ppm dinotefuran, and pollen paste with 5.65 ppm dinotefuran, respectively. Dinotefuran is denoted by DF, and 10 ppm dinotefuran is equivalent to 10% of the recommended concentration (100 ppm) for exterminating stink bugs.

### Methods used in field experiments

In order to enhance the accuracy of experiment, it can be considered to increase the number of experimental colonies from the statistical point of view and to improve the accuracy of measurement in each experiment. In this study, we have tried to improve the accuracy of measurement as follows:

The numbers of honeybees and capped brood in a hive are often estimated from the weight of a hive and the area ratio on each comb occupied by sealed cells, respectively. These are a simple and easy method which can roughly grasp the change of the number of each member in a honeybee colony but may lack accuracy. For example, in the estimation of honeybee number by the weight method the weight of an individual honeybee is incomparably lighter than the whole weight of a hive and the instrumental errors in measurement can be equivalent to the weight of a few hundred honeybees. And it is hard to differentiate the weight of honeybees from that of the others such as honey, bee-bread, comb, propolis etc in a hive which change with time. The estimation of the number of capped brood by the area method may include the difficulty when the area is occupied heterogeneously by dotted capped brood and the others. We have tried to accurately count honeybees and capped brood one by one on the photos of combs and the inside of a hive. Though we have tried to develop a new automatic counting software system with the operation of binarizing photo images of combs and the inside of a hive, we cannot succeed in accurate counting of them because exposure conditions to take photos are unstable in the field. The system cannot always accurately count overlaid bees, ones on a blurred image, ones on a low or an uneven contrast image and/or ones on a low brightness image, and capped brood differentiating from sealed honey in sealed cells even when the threshold is changed. Therefore, we have tried to count honeybees and capped brood directly and patiently one by one with a hand-operated counter judging visually to aim for accuracy.

Five hives, in each of which three combs numbered and ordered numerically and a feeder were installed, were sited on a hill facing east and being aligned north-south. The total number of adult bees on all combs and a feeder and the inside of a hive box (4 walls and the bottom) was counted directly from photographs (sometimes enlarged) of them with a counter. The total number of capped brood was counted in a similar manner, after shaking the bees off each comb. The total number of dead bees in and around a hive were counted, which was placed on a large tray, one by one with a pair of tweezers. Consumption of foods (sugar syrup and pollen paste) by the honeybees was accurately measured with a weighing instrument after removing dead bees. A queen in a hive was photographically recorded on each observation date, as were specific situations such as the presence of chalk brood or wax moth larvae and Asian giant hornet attacks. During the experimental period, the state of a hive was recorded with a digital camera at intervals of 1 h.

Experiments started in July, after the swarming season, and were performed early in the morning on fine or cloudy days, before the foraging bees left a hive. They were performed in an apiary where honeybees could freely visit flowers in the field, thus allowing to avoid consumption of the food provided (sugar syrup and pollen paste containing a pesticide) in the case that the pesticide was repellent to honeybees. Both new foods (sugar syrup and pollen paste), which were prepared according to experimental conditions, were fed to a hive after weighing old ones which were removed from the hive every observation date. In DF-Low/Syrup and DF-Low/Pollen, the pesticide (dinotefuran) continued to be administered into a colony till extinction and just before wintering, respectively. On the other hand, in DF-High/Syrup and DF-High/Pollen, the pesticide was administered only for the first time around (for the first 7 days). The surviving colonies escaped from a collapse appeared vigorous before wintering. Experiments were recorded by still and moving photography, and the results could be accessed at any time, if necessary.

## Results

### Long-term observations

Table 2 shows our observational results in this work. General information from these observations can give an overview of changes in experimental situations and conditions, such as the numbers of combs added later as occasion arises, the incidences of chalk brood, or wax moth mites, and attacks by Asian giant hornets and the number of dead bees.

**Table 2.** Memorandum of observational results in this work

| Date | Days | Control |  | DF-Low/Syrup |  | DF-Low/Pollen |  | DF-High/Syrup |  | DF-High/Pollen |  |
| --- | --- | --- | --- | --- | --- | --- | --- | --- | --- | --- | --- |
|  |  | Combs | Note | Combs | Note | Combs | Note | Combs | Note | Combs | Note |
| 9-Jul-11 | 0 | 3 | No dead bees: No mites | 3 | No dead bees: No mites | 3 | No dead bees: No mites | 3 | No dead bees: No mites | 3 | No dead bees: No mites |
| 10-Jul-11 | 0 | 3 | Without the check in the inside of hive<br>No dead bees | 3 | Without the check in the inside of hive<br>No dead bees | 3 | Without the check in the inside of hive<br>No dead bees | 3 | Without the check in the inside of hive<br>Many dead bees around the hive (only by the appearance without counting) | 3 | Without the check in the inside of hive<br>Many dead bees around the hive (only by the appearance without counting) |
| 16-Jul-11 | 7 | 3 | 2 dead bees: No mites | 3 | 444 dead bees: No mites | 3 | 113 dead bees: No mites | 3 | Stop of the pesticide administration<br>Many (3314) dead bees around and inside the hive: A few mites | 3 | Stop of the pesticide administration<br>Many (1899) dead bees around and inside the hive: No mites<br>Queen-loss (she died in the hive) |
| 22-Jul-11 | 13 | 4 | Addition of a comb: No dead bees<br>No mites | 3 | 137 dead bees: No mites | 3 | 113 dead bees: No mites | 3 | 7 dead bees: No mites | 3 | 319 dead bees: No mites |
| 29-Jul-11 | 20 | 5 | Addition of a comb: 5 dead bees<br>No mites | 3 | 279 dead bees: No mites | 3 | 110 dead bees: No mites | 3 | 130 dead bees: No mites | 3 | 196 dead bees: 3 mites<br>1 wax-moth larvae |
| 6-Aug-11 | 28 | 5 | No dead bees: No mites | 3 | 152 dead bees: No mites | 3 | 45 dead bees | 3 | 80 dead bees: No mites | 3 | Termination of the experiment<br>2 dead bees: No mites:<br>1 wax-mph larva: Stock of foods |
| 12-Aug-11 | 34 | 6 | Addition of a comb: Installation of a hornet-capture: 4 dead bees: No mites | 3 | Installation of a hornet-capture<br>157 dead bees: No mites | 3 | Installation of a hornet-capture<br>112 dead bees: No mites | 3 | Installation of a hornet-capture<br>88 dead bees: No mites |  |  |
| 18-Aug-11 | 40 | 6 | 12 dead bees: No mites | 3 | 59 dead bees: No mites | 3 | 115 dead bees: No mites | 3 | 69 dead bees: No mites |  |  |
| 26-Aug-11 | 48 | 6 | 13 dead bees: No mites | 3 | A feeble queen: 160 dead bees<br>No mites | 3 | 116 dead bees: No mites | 3 | 10 dead bees: No mites |  |  |
| 10-Sep-11 | 63 | 6 | 14 dead bees: No mites | 3 | 115 dead bees: No mites | 3 | 144 dead bees: No mites | 3 | 42 dead bees: No mites |  |  |
| 17-Sep-11 | 70 | 6 | 23 dead bees: No mites | 3 | 15 dead bees: No mites | 3 | 112 dead bees: No mites | 3 | 134 dead bees: No mites |  |  |
| 24-Sep-11 | 77 | 6 | Attack by hornets: 86 dead bees<br>No mites | 3 | Some dead brood: 7 dead bees<br>No mites | 3 | 115 dead bees: No mites | 3 | Attack by hornets: 583 dead bees<br>No mites |  |  |
| 29-Sep-11 | 82 | 6 | 6 dead bees: No mites | 3 | Some dead brood & a few wax-moth larvae larvae : 6 dead bees: No mites | 3 | 69 dead bees: No mites | 3 | 48 dead bees: No mites |  |  |
| 7-Oct-11 | 90 | 6 | 4 dead bees: No mites | 3 | A few wax-moth larvae & some hive beetles: 9 dead bees: No mites | 3 | 92 dead bees: No mites | 3 | 13 dead bees: No mites |  |  |
| 21-Oct-11 | 104 | 6 | 2 dead bees: No mites | 3 | Extinction & many wax-moth larvae (Termination of the experiment)<br>1 dead bee: No mites: Stock of foods | 3 | 98 dead bees: No mites | 3 | 5 dead bees: No mites |  |  |
| 30-Oct-11 | 113 | 6 | 12 dead bees: No mites |  |  | 3 | 15 dead bees: No mites | 3 | 3 dead bees: No mites |  |  |
| 4-Nov-11 | 118 | 6 | 5 dead bees: No mites |  |  | 3 | 87 dead bees: 1 mite | 3 | 1 dead bees: No mites |  |  |
| 18-Nov-11 | 132 | 6 | 35 dead bees: No mites |  |  | 3 | 47 dead bees: No mites | 3 | Queen-loss: 24 dead bees: No mites |  |  |
| 26-Nov-11 | 140 | 6 | 305 dead bees: No mites |  |  | 3 | Attack by hornets: 316 dead bees<br>No mites | 3 | 59 dead bees: No mites |  |  |
| 3-Dec-11 | 148 | 6 | Attack by hornets: 113 dead bees:<br>No mites |  |  | 3 | Attack by hornets: 119 dead bees<br>After the stop of pollen paste with dinotefuran and sugar syrup without dinotefuran kept being fed. No mites | 3 | (Termination of the experiment)<br>22 dead bees: No mites<br>Stock of foods |  |  |
| 17-Dec-11 | 162 | 6 | Stop of sugar syrup feeding and start of wintering |  |  | 3 | Stop of sugar syrup feeding and start of wintering |  |  |  |  |
| 16-Feb-12 | 223 | 6 | Wintering |  |  | 3 | Extinction (Failure in wintering)<br>(Termination of the experiment)<br>Stock of foods |  |  |  |  |
| 2-Apr-12 | 269 | 6 | Success in wintering: 1 mite |  |  | 3 |  |  |  |  |  |

(Note) Total number of dead bees in the hive at the observational date = Number of dead bees near the hive + Number of dead bees inside the hive

It has been revealed that high-concentration pesticides caused almost instant deaths of many bees and that few mites were present in every hive during the experimental period even after very few adults remained. Almost no wax moth larvae were present except in DF-Low/Syrup where they began to appear about three weeks before the extinction and greatly increased just before the extinction. Each queen continued to survive to extinction of the colony except for the case of DF-High/Pollen.

### Measurement of number of adult bees, capped brood, and dead bees

Table 3 shows the numbers of adult bees, capped brood, and dead bees in this work.

**Table 3.** The numbers of adult bees, capped brood & dead bees

| Date | Days | Control<br>pesticide-free |  |  | DF-Low/Syrup<br>1 ppm of dinotefuran |  |  | DF-Low/Pollen<br>0.565 ppm of dinotefuran |  |  | DF-High/Syrup<br>10 ppm of dinotefuran |  |  | DF-High/Pollen<br>5.65 ppm of dinotefuran |  |  |
| --- | --- | --- | --- | --- | --- | --- | --- | --- | --- | --- | --- | --- | --- | --- | --- | --- |
|  |  | Adult | Brood | Dead | Adult | Brood | Dead | Adult | Brood | Dead | Adult | Brood | Dead | Adult | Brood | Dead |
| 9-Jul-11 | 0 | 3392 | 5819 | 0 | **2965** | **4263** | **0** | **2158** | **2556** | **0** | **3295** | **3880** | **0** | **1659** | **6093** | **0** |
| 16-Jul-11 | 7 | 6827 | 3644 | 2 | **4257** | **1287** | **444** | **3755** | **2377** | **113** | **1102** | **986** | **3314** | **[430]** | **[2865]** | **[1899]** |
| 22-Jul-11 | 13 | 6399 | 4692 | 0 | **5233** | **1655** | **137** | **3493** | **3207** | **113** | 1407 | 232 | 7 | [330] | [1165] | [319] |
| 29-Jul-11 | 20 | 7737 | 4861 | 5 | **3131** | **1034** | **279** | **4025** | **2743** | **110** | 1451 | 939 | 130 | [17] | [790] | [196] |
| 6-Aug-11 | 28 | 7893 | 7179 | 0 | **2393** | **1591** | **152** | **4372** | **3480** | **45** | 1384 | 2370 | 80 | [1] | [729] | [2] |
| 12-Aug-11 | 34 | 7675 | 8390 | 4 | **1762** | **2288** | **157** | **4513** | **3217** | **112** | 1752 | 3217 | 88 |  |  |  |
| 18-Aug-11 | 40 | 8873 | 6125 | 12 | **2298** | **1137** | **59** | **4829** | **1933** | **115** | 2257 | 946 | 69 |  |  |  |
| 26-Aug-11 | 48 | 9327 | 5797 | 13 | **1623** | **1757** | **160** | **4276** | **2099** | **116** | 2271 | 913 | 10 |  |  |  |
| 10-Sep-11 | 63 | 9249 | 8803 | 14 | **1253** | **318** | **115** | **4620** | **1343** | **144** | 2096 | 545 | 42 |  |  |  |
| 17-Sep-11 | 70 | 9762 | 8327 | 23 | **997** | **341** | **15** | **4241** | **1546** | **112** | 1760 | 707 | 134 |  |  |  |
| 24-Sep-11 | 77 | 11252 | 7034 | 86 | **462** | **453** | **7** | **3862** | **2693** | **115** | 908 | 772 | 583 |  |  |  |
| 29-Sep-11 | 82 | 10736 | 5810 | 6 | **342** | **230** | **6** | **3792** | **2445** | **69** | 996 | 505 | 48 |  |  |  |
| 7-Oct-11 | 90 | 12015 | 6100 | 4 | **169** | **109** | **9** | **4232** | **2082** | **92** | 1076 | 12 | 13 |  |  |  |
| 21-Oct-11 | 104 | 11253 | 4100 | 2 | **[0]** | **[45]** | **[1]** | **4225** | **1935** | **98** | 860 | 220 | 5 |  |  |  |
| 30-Oct-11 | 113 | 10958 | 4300 | 12 |  |  |  | **4044** | **2105** | **15** | 895 | 147 | 3 |  |  |  |
| 4-Nov-11 | 118 | 10654 | 3900 | 5 |  |  |  | **3675** | **2665** | **87** | 790 | 245 | 1 |  |  |  |
| 18-Nov-11 | 132 | 11303 | 3430 | 35 |  |  |  | **4022** | **1720** | **47** | [779] | [160] | [24] |  |  |  |
| 26-Nov-11 | 140 | 12390 | 1750 | 305 |  |  |  | **3806** | **1034** | **316** | [630] | [48] | [59] |  |  |  |
| 3-Dec-11 | 148 | 12109 | 2038 | 113 |  |  |  | **2992** | **840** | **119** | [393] | [21] | [22] |  |  |  |
| 17-Dec-11 | 162 | 11811 | 212 |  |  |  |  | 2433 | 157 |  | [0] | [0] |  |  |  |  |
| 16-Feb-12 | 223 | 10514 | 0 |  |  |  |  | [0] | [0] |  |  |  |  |  |  |  |
| 2-Apr-12 | 269 | 9622 | 1873 |  |  |  |  |  |  |  |  |  |  |  |  |  |

Total number of dead bees at each observation date = Number of dead bees near the hive + Number of dead bees inside the hive
The "**bold red**" figures denote a state that foods (sugar syrup, pollen paste) with a pesticide (dinotefuran) were fed into a colony and the "Times New Roman" ones denote pesticide-free foods. Foods with a pesticide were changed to pesticide-free ones in the morning of measurement date: E.g., in DF-High/Syrup and DF-High/Pollen, the change was made in the morning of July 16th. The [bracketed figures] denote a state that the queen had been lost.

Figures 1–3 show changes in the number of adult bees, capped brood and dead bees, respectively. Administration of the pesticide led to the decreases in adult bees and capped brood as shown in Figures 1 and 2. Experimental colonies of DF-Low/Syrup, DF-Low/Pollen, and DF-High/Syrup dwindled away to nothing through the aspects of a CCD (the existence of a queen with some bees and capped brood, stock of foods, few dead bees around a hive) except for DF-High/Pollen. Such aspects of a CCD has been already assumed similarly in our previous work (Yamada *et al.*, 2012) conducted in 2010. A photographic image of a CCD is shown in Figure 4. On the other hand, in DF-High/Pollen a queen was dead before the second administration of the pesticide.

**Figure 1.**
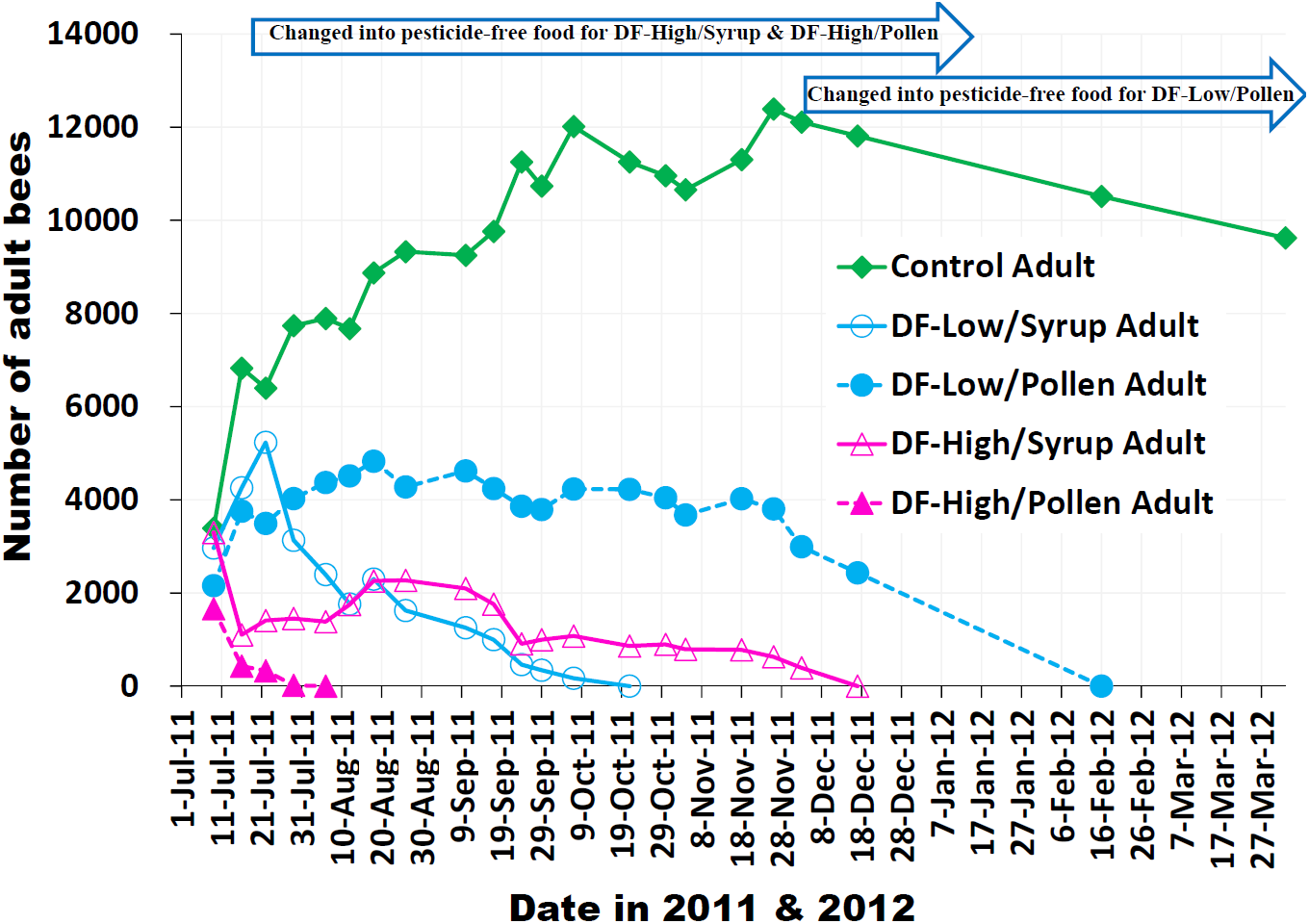
Change in number of Adult bees The pesticide (dinotefuran) was administered through either sugar syrup or pollen paste, the latter consisting of 10 parts of pollen without dinotefuran and 13 parts of sugar syrup with dinotefuran. This was prepared by mixing the pollen substitute BB Food A^®^ (Bee Culture Laboratory Co., Japan) with pure pollen at a ratio of 1:1. Control: without dinotefuran. DF-Low/Syrup: 1 ppm dinotefuran in sugar syrup. DF-Low/Pollen: 0.565 ppm dinotefuran in pollen paste, consisting of 10 parts of pollen without dinotefuran and 13 parts of sugar syrup with 1 ppm dinotefuran. DF-High/Syrup: 10 ppm dinotefuran in sugar syrup; the sugar syrup without dinotefuran was administered on and after July 16, 2011. DF-High/Pollen: 5.65 ppm dinotefuran in pollen paste, consisting of 10 parts of pollen without dinotefuran and 13 parts of sugar syrup with 10 ppm dinotefuran; the pollen paste without dinotefuran was administered on and after July 16, 2011. In DF-Low/Pollen, the administration of dinotefuran through pollen paste was discontinues with keeping feeding pesticide-free sugar syrup on December 3, 2011. On December 17, 2011, colonies of Control and DF-Low/Pollen entered their wintering after pesticide-free sugar syrup was discontinued. We observed the two colonies during wintering on February 16, 2012 choosing a fine day in order to avoid an adverse effect on the colonies and found that the colony of DF-Low/Pollen had become extinct, whereas the colony of Control survived.

**Figure 2.**
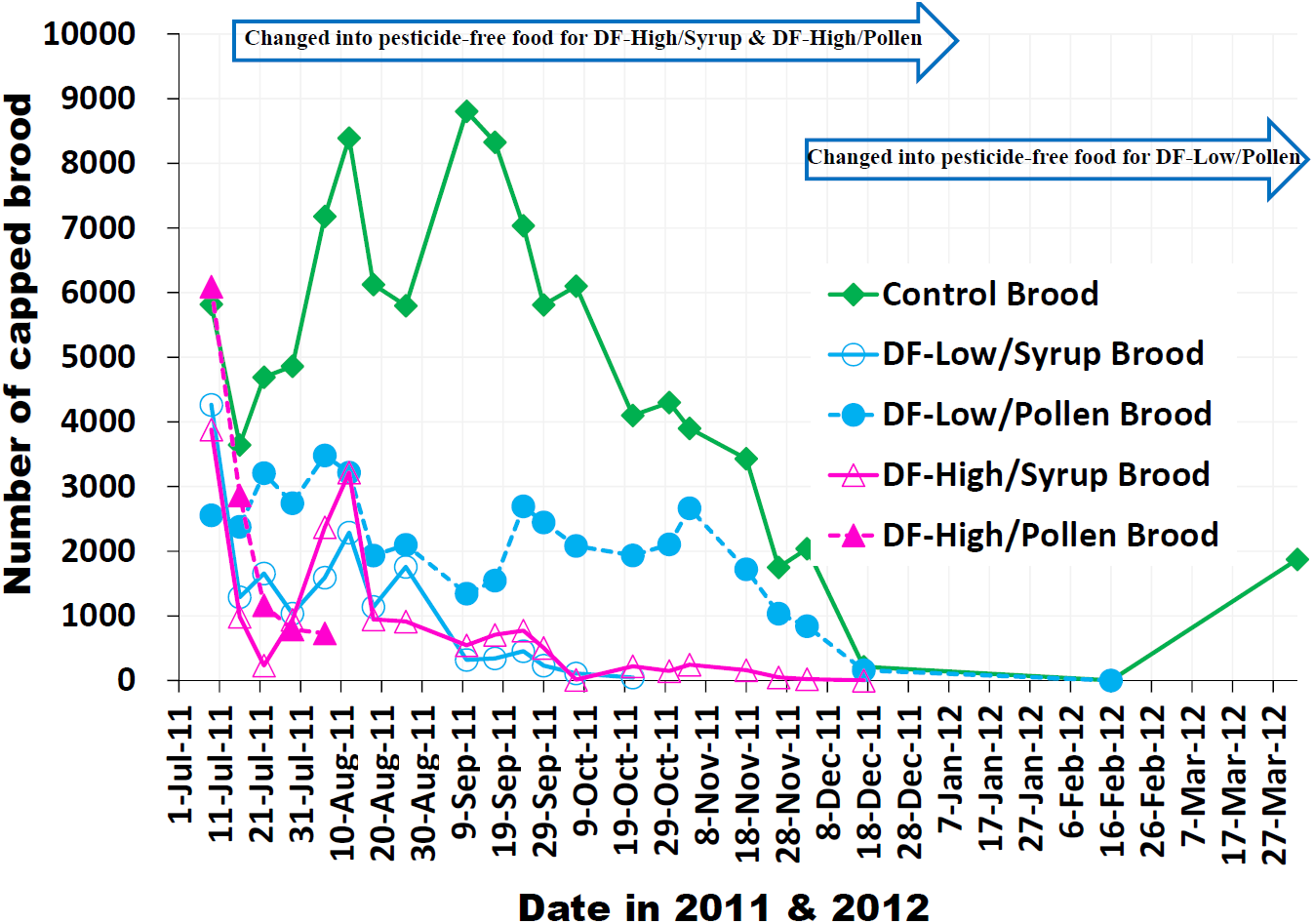
Change in number of capped brood

**Figure 3.**
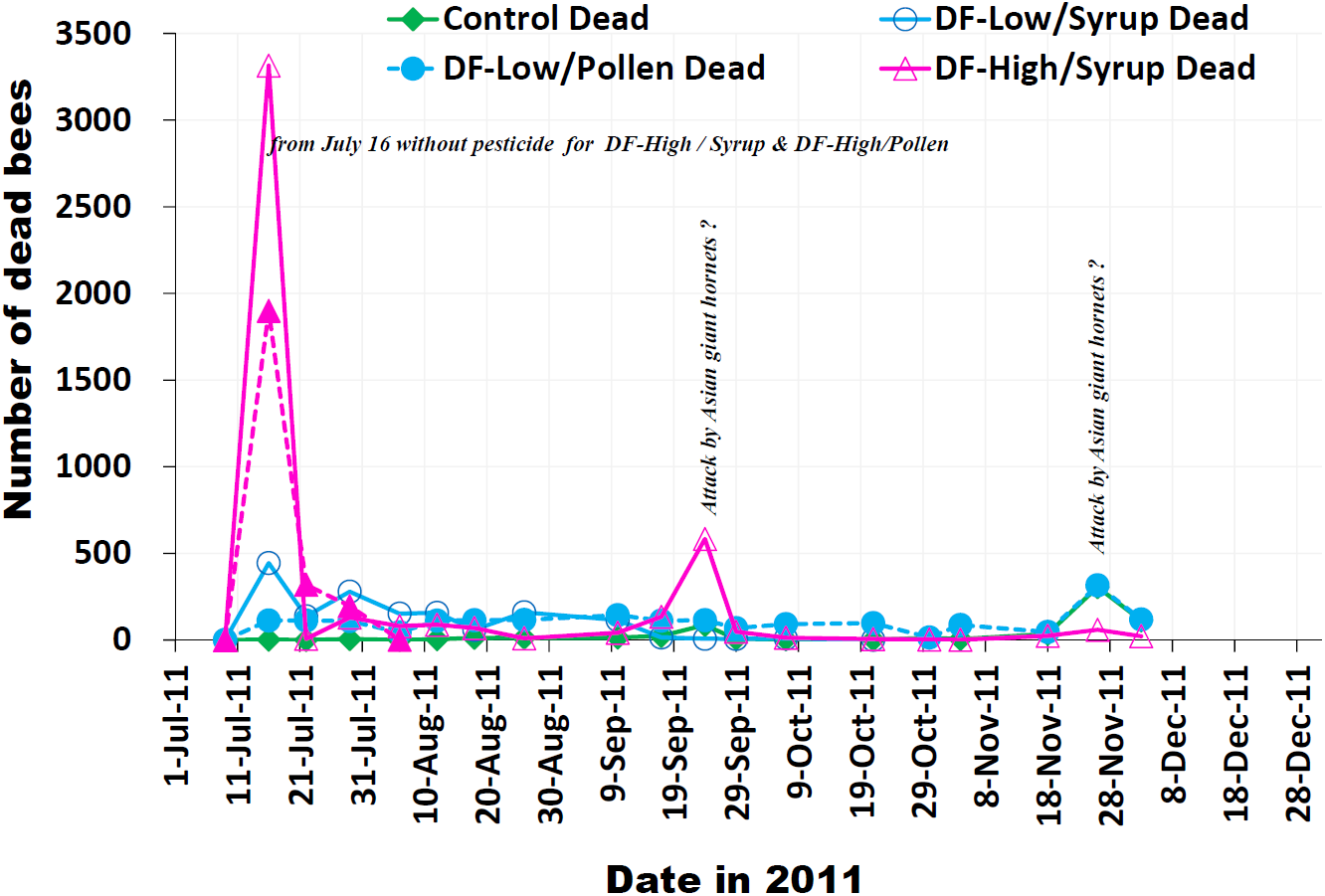
Change in number of dead bees The marked increase in the number of dead bees observed immediately after the start of the experiment was because of pesticide (dinotefuran) toxicity. Incidentally, from mid-September, the sudden increases of dead bee was due to attack by Asian giant hornets.

Table 3 and Figures 1–3 suggest the followings under the support of the records with digital cameras in front of hives at intervals of 1 h:

In DF-Low/Syrup where sugar syrup containing 1 ppm of dinotefuran continued to be fed to a colony till extinction, adult bees and capped brood gradually decreased in number under the existence of a queen and far fewer dead bees than the decrease of adult bees and finally the colony become extinct. In DF-Low/Pollen where pollen paste containing 0.565 ppm of dinotefuran continued to be fed to a colony till the morning of November 26^th^, 2011 (just before wintering) and after dinotefuran-free one was fed till extinction, adult bees and capped brood did not much change in number till the start of wintering under the existence of a queen and as few dead bees as the ordinary level and they became zero during wintering. In DF-Low/Syrup and DF-Low/Pollen, a queen survived to colony extinction, with a gradual decrease in the numbers of adult bees and capped brood, and the presence of as few dead bees as the ordinary level giving the appearance of a CCD prior to extinction.

In DF-High/Syrup where sugar syrup containing 10 ppm of dinotefuran was fed to a colony for first seven days (only once) and after that pesticide-free one continued to be fed, adult bees and capped brood sharply decreased in number with many dead bees just after administration of the pesticide and they gradually decreased in number under the existence of a queen and few dead bees and finally they become zero. In DF-High/Pollen, pollen paste containing 5.65 ppm of dinotefuran was fed to a colony for first seven days (only once) with a queen being dead in the meantime, and after that pesticide-free one continued to be fed. Adult bees and capped brood sharply decreased in number with many dead bees without a queen and they decreased in number with some dead bees and finally the colony become extinct.

In DF-High/Syrup and DF-High/Pollen, mass dead bees were found in front of a hive on the day after administration of the pesticide as described in Table 2. In DF-High/Syrup, a queen survived to colony extinction, with sharp decreases in the numbers of adult bees and capped brood just after the administration of dinotefuran, followed by their gradual decreases with the colony giving the appearance of a CCD where a queen, brood and a small number of workers exist with few dead bees near a hive and some bee-foods (honey, bee-bread). On the other hand, in DF-High/Pollen, a queen was lost immediately after the administration of dinotefuran, and there were many dead bees present. This was followed by a sharp decrease in the number of adult bees and capped brood and the colony rapidly become extinct. These results suggest that when a pesticide such as dinotefuran is administered through pollen paste, it may impact a queen more strongly than when it is administered through sugar syrup.

Figure 3 indicates that the number of dead bees remained almost constant at a low level, except for the periods immediately after the administration of high-concentration dinotefuran and when attack by Asian giant hornets was recorded.

Colonies of DF-Low/Syrup, DF-High/Syrup and DF-High/Pollen had already collapsed before wintering. Although the surviving DF-Low/Pollen colony appeared active just before wintering on December 3, whose number of adult bees on that date was greater than that at the start of the experiment on July 9, the colony was confirmed to be unsuccessful in wintering by mid-February the following year, as shown in Figure 5. This seems to be because the administration of the pesticide through pollen mainly affects brood and newborn bees before winter and makes the longevity of adult bees which have eclosed from the brood in DF-Low/Pollen shorter than that in Control. The control colony succeeded in wintering and was vigorous after the end of experiment (April 2^nd^, 2012).

**Figure 4.**
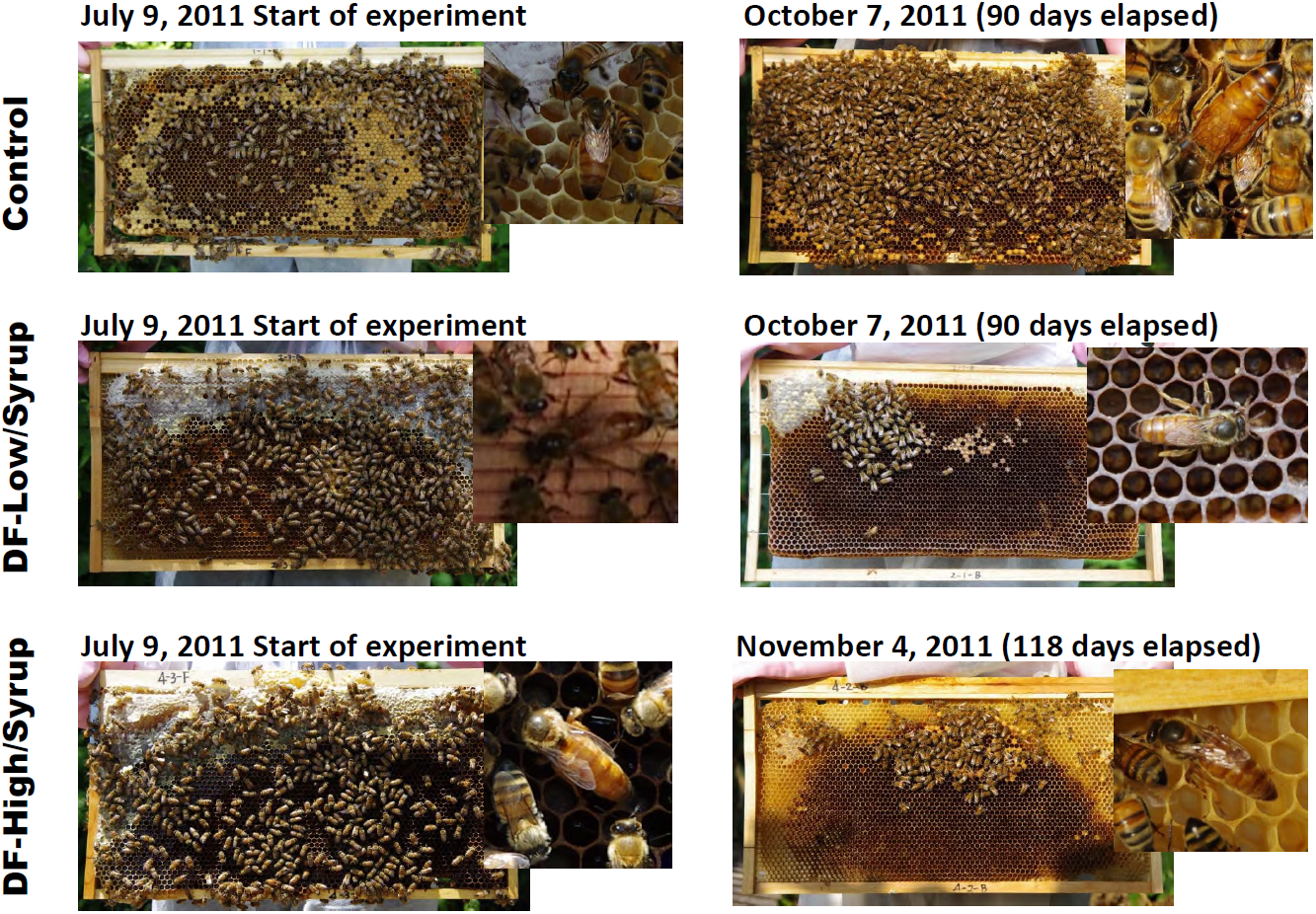
Difference in state on comb between at the start of experiment and just before colony extinction Administration of the pesticide (dinotefuran) in DF-Low/Syrup was continued to colony extinction, but only at the start of the experiment in DF-High/Syrup. A queen remained alive on the verge of colony extinction, when the previously vigorous colony assumed the appearance of a CCD.

**Figure 5.**
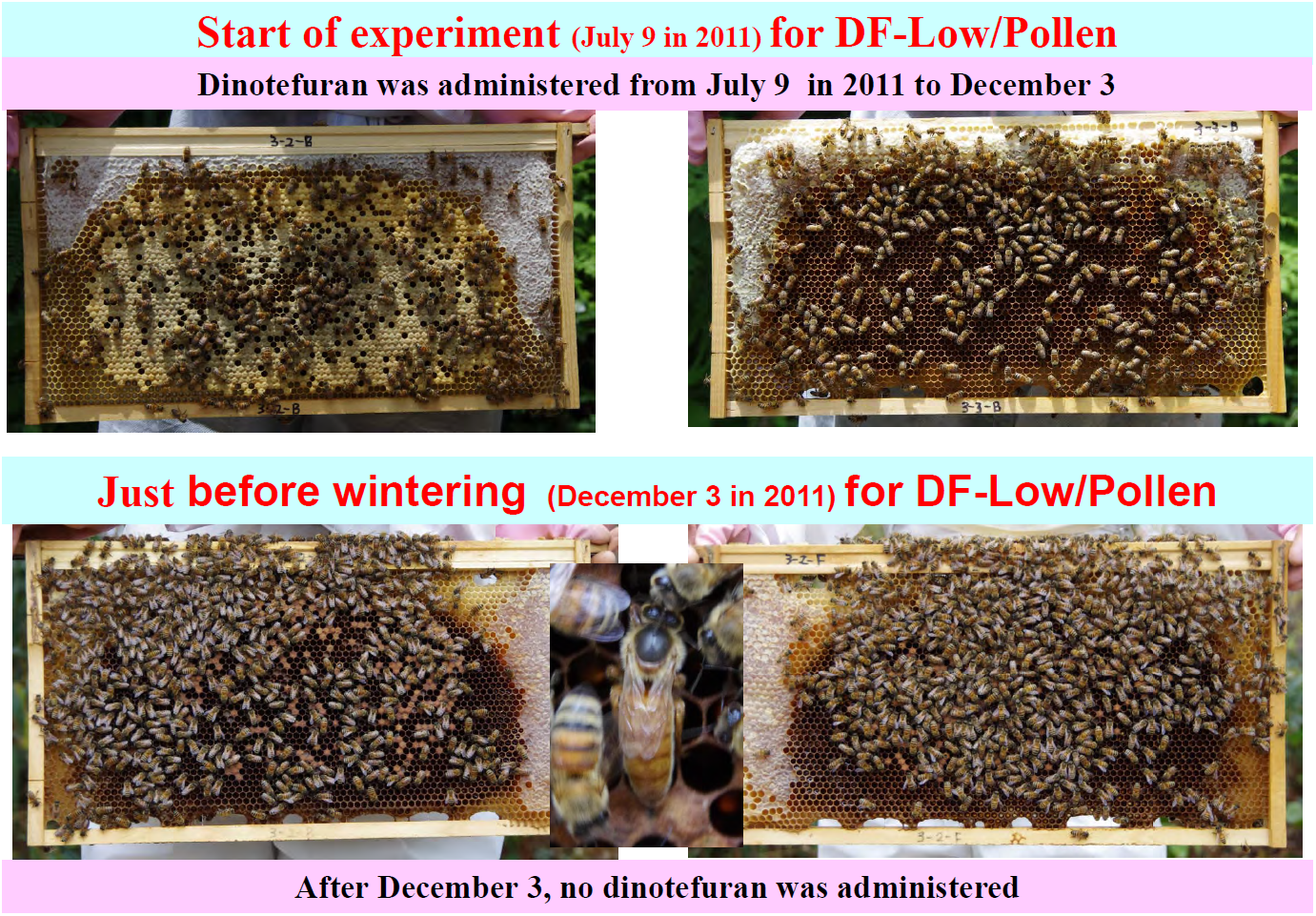
Comparison of the DF-Low/Pollen colony at the start of the experiment and just before wintering The upper and lower images are representative combs with honeybees at the start of the experiment and immediately before wintering (existence of the queen confirmed), respectively. The colony appeared vigorous before wintering but became extinct during wintering.

### Intake of pesticide in colonies

Table 4 shows the change in the consumption of foods (sugar syrup, pollen paste) and the change in the intake of dinotefuran (pesticide) taken from foods in this work. Figure 6 shows the change in the intake of dinotefuran for experimental colonies.

**Figure 6.**
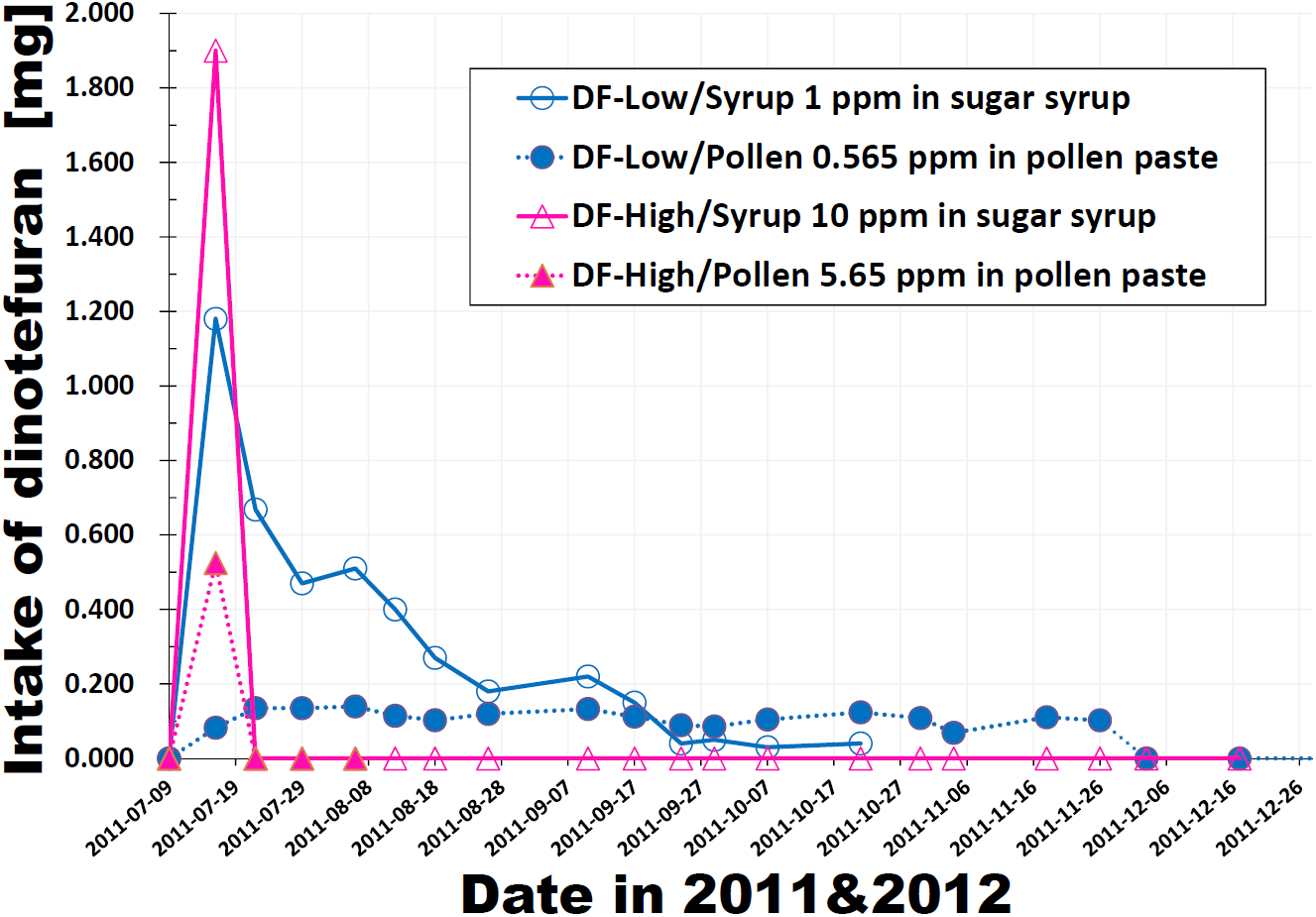
Change in the total intake of dinotefuran In DF-High/Sugar syrup and In DF-High/Pollen, the pesticide (dinotefuran) was administered only once at the start of the experiment.

**Table 4.** Change in the intakes of foods (sugar syrup, pollen paste) and pesticide (dinotefuran) taken from foods

| Date | Elapsed days | Control |  |  |  | DF-Low/Syrup |  |  |  |  |  | DF-Low/Pollen |  |  |  |  |  | DF-High/Syrup |  |  |  |  |  | DF-High/Pollen |  |  |  |  |  |
| --- | --- | --- | --- | --- | --- | --- | --- | --- | --- | --- | --- | --- | --- | --- | --- | --- | --- | --- | --- | --- | --- | --- | --- | --- | --- | --- | --- | --- | --- |
|  |  | Pesticide (dinotefuran) - free |  |  |  | 1 ppm of dinotefuran in sugar syrup |  |  |  |  |  | 0.565 ppm of dinotefuran in pollen paste |  |  |  |  |  | 10 ppm of dinotefuran in sugar syrup |  |  |  |  |  | 5.65 ppm of dinotefuran in pollen paste |  |  |  |  |  |
|  |  | Amount of consumption |  |  |  | Amount of consumption |  |  |  |  |  | Amount of consumption |  |  |  |  |  | Amount of consumption |  |  |  |  |  | Amount of consumption |  |  |  |  |  |
|  |  | Sugar syrup [g] |  | Pollen paste [g] |  | Sugar syrup [g] |  | Pollen paste [g] |  | Dinotefuran [mg] |  | Sugar syrup [g] |  | Pollen paste [g] |  | Dinotefuran [mg] |  | Sugar syrup [g] |  | Pollen paste [g] |  | Dinotefuran [mg] |  | Sugar syrup [g] |  | Pollen paste [g] |  | Dinotefuran [mg] |  |
|  |  | Interval | Integrated | Interval | Integrated | Interval | Integrated | Interval | Integrated | Interval | Integrated | Interval | Integrated | Interval | Integrated | Interval | Integrated | Interval | Integrated | Interval | Integrated | Interval | Integrated | Interval | Integrated | Interval | Integrated | Interval | Integrated |
| 9-Jul-11 | 0 | 0 | 0 | 0 | 0 | 0 | 0 | 0 | 0 | 0.000 | 0.000 | 0 | 0 | 0 | 0 | 0.0000 | 0.0000 | 0 | 0 | 0 | 0 | 0.000 | 0.000 | 0 | 0 | 0 | 0 | 0.000 | 0.000 |
| 16-Jul-11 | 7 | 1500 | 1500 | 250 | 250 | 1180 | 1180 | 180 | 180 | 1.180 | 1.180 | 1222 | 1222 | 146 | 146 | 0.0825 | 0.0825 | 190 | 190 | 161 | 161 | 1.900 | 1.900 | [90] | [90] | [93] | [93] | [0.525] | [0.525] |
| 22-Jul-11 | 13 | 1500 | 3000 | 250 | 500 | 668 | 1848 | 165 | 345 | 0.668 | 1.848 | 1020 | 2242 | 239 | 385 | 0.1350 | 0.2175 | 90 | 280 | 113 | 274 | 0.000 | 1.900 | [220] | [310] | [87] | [180] | [0] | [0.525] |
| 29-Jul-11 | 20 | 1500 | 4500 | 250 | 750 | 470 | 2318 | 194 | 539 | 0.470 | 2.318 | 1270 | 3512 | 239 | 624 | 0.1350 | 0.3525 | 1010 | 1290 | 164 | 438 | 0.000 | 1.900 | [820] | [1130] | [53] | [233] | [0] | [0.525] |
| 6-Aug-11 | 28 | 1500 | 6000 | 216 | 966 | 510 | 2828 | 206 | 745 | 0.510 | 2.828 | 1500 | 5012 | 247 | 871 | 0.1396 | 0.4921 | 370 | 1660 | 158 | 596 | 0.000 | 1.900 | [0] | [1130] | [0] | [233] | [0] | [0.525] |
| 12-Aug-11 | 34 | 1500 | 7500 | 215 | 1181 | 400 | 3228 | 144 | 889 | 0.400 | 3.228 | 810 | 5822 | 203 | 1074 | 0.1147 | 0.6068 | 220 | 1880 | 143 | 739 | 0.000 | 1.900 |  |  |  |  |  |  |
| 18-Aug-11 | 40 | 1500 | 9000 | 224 | 1405 | 270 | 3498 | 122 | 1011 | 0.270 | 3.498 | 1000 | 6822 | 182 | 1256 | 0.1028 | 0.7096 | 170 | 2050 | 105 | 844 | 0.000 | 1.900 |  |  |  |  |  |  |
| 26-Aug-11 | 48 | 1500 | 10500 | 248 | 1653 | 180 | 3678 | 157 | 1168 | 0.180 | 3.678 | 1120 | 7942 | 212 | 1468 | 0.1198 | 0.8294 | 220 | 2270 | 106 | 950 | 0.000 | 1.900 |  |  |  |  |  |  |
| 10-Sep-11 | 63 | 1500 | 12000 | 250 | 1903 | 220 | 3898 | 116 | 1284 | 0.220 | 3.898 | 910 | 8852 | 235 | 1703 | 0.1328 | 0.9622 | 280 | 2550 | 114 | 1064 | 0.000 | 1.900 |  |  |  |  |  |  |
| 17-Sep-11 | 70 | 1500 | 13500 | 236 | 2139 | 150 | 4048 | 86 | 1370 | 0.150 | 4.048 | 1090 | 9942 | 199 | 1902 | 0.1124 | 1.0746 | 380 | 2930 | 82 | 1146 | 0.000 | 1.900 |  |  |  |  |  |  |
| 24-Sep-11 | 77 | 1500 | 15000 | 226 | 2365 | 40 | 4088 | 48 | 1418 | 0.040 | 4.088 | 470 | 10412 | 158 | 2060 | 0.0893 | 1.1639 | 80 | 3010 | 69 | 1215 | 0.000 | 1.900 |  |  |  |  |  |  |
| 29-Sep-11 | 82 | 1500 | 16500 | 230 | 2595 | 50 | 4138 | 20 | 1438 | 0.050 | 4.138 | 360 | 10772 | 152 | 2212 | 0.0859 | 1.2498 | 40 | 3050 | 30 | 1245 | 0.000 | 1.900 |  |  |  |  |  |  |
| 7-Oct-11 | 90 | 1500 | 18000 | 211 | 2806 | 30 | 4168 | 11 | 1449 | 0.030 | 4.168 | 610 | 11382 | 185 | 2397 | 0.1045 | 1.3543 | 60 | 3110 | 46 | 1291 | 0.000 | 1.900 |  |  |  |  |  |  |
| 21-Oct-11 | 104 | 1500 | 19500 | 250 | 3056 | [40] | [4208] | [10] | [1459] | [0.040] | [4.208] | 1150 | 12532 | 219 | 2616 | 0.1237 | 1.4780 | 40 | 3150 | 70 | 1361 | 0.000 | 1.900 |  |  |  |  |  |  |
| 30-Oct-11 | 113 | 1500 | 21000 | 250 | 3306 |  |  |  |  |  |  | 1500 | 14032 | 192 | 2808 | 0.1085 | 1.5865 | 80 | 3230 | 33 | 1394 | 0.000 | 1.900 |  |  |  |  |  |  |
| 4-Nov-11 | 118 | 1500 | 22500 | 219 | 3522 |  |  |  |  |  |  | 1120 | 15152 | 122 | 2930 | 0.0689 | 1.6554 | 200 | 3430 | 20 | 1414 | 0.000 | 1.900 |  |  |  |  |  |  |
| 18-Nov-11 | 132 | 1500 | 24000 | 240 | 3762 |  |  |  |  |  |  | 1500 | 16652 | 196 | 3126 | 0.1107 | 1.7661 | [170] | [3600] | [34] | [1448] | [0] | [1.900] |  |  |  |  |  |  |
| 26-Nov-11 | 140 | 1330 | 25330 | 239 | 4001 |  |  |  |  |  |  | 860 | 17512 | 181 | 3307 | 0.1023 | 1.8684 | [50] | [3650] | [10] | [1458] | [0] | [1.900] |  |  |  |  |  |  |
| 3-Dec-11 | 148 | 1500 | 26830 | 242 | 4243 |  |  |  |  |  |  | 350 | 17862 | 123 | 3430 | 0.0000 | 1.8684 | [10] | [3660] | [10] | [1468] | [0] | [1.900] |  |  |  |  |  |  |
| 17-Dec-11 | 162 | 0 | 26830 | 0 | 4243 |  |  |  |  |  |  | 290 | 18152 | 0 | 3430 | 0.0000 | 1.8684 | [0] | [3660] | [0] | [1468] | [0] | [1.900] |  |  |  |  |  |  |
| 16-Feb-12 | 223 | 0 | 26830 | 0 | 4243 |  |  |  |  |  |  | [0] | [18152] | [0] | [3430] | [0] | [1.8684] |  |  |  |  |  |  |  |  |  |  |  |  |
| 2-Apr-12 | 269 | 0 | 26830 | 0 | 4243 |  |  |  |  |  |  |  |  |  |  |  |  |  |  |  |  |  |  |  |  |  |  |  |  |

The "**bold red**" figures denote a state that foods (sugar syrup, pollen paste) with a pesticide (dinotefuran) were fed into a colony and the "Times New Roman" ones denote pesticide-free foods.
The bracketed [ ] figures denote a state that the queen had been lost.

Table 4 and Figure 6 indicate that high-concentration pesticide was mostly taken by honeybees for a short time after the start of the experiment but that low-concentration pesticide was taken over a longer period before colony extinction. The total intake of pesticide in a colony can involve honey or bee bread stored in combs as noxious provisions

### Grand total number of honeybees having taken pesticide in an experimental colony

Here, we consider two cases on a period when honeybees were exposed to a pesticide in an experimental colony: CASE 1 is the shortest period case where the pesticide was immediately ingested by honeybees soon after administered and the period is equal to the administration one; CASE 2 is the longest one where the pesticide was partly stored in cells and continued to be ingested by honeybees till the colony extinction. In a control colony the period to obtain the grand total number is the feeding period of sugar syrup or pollen paste.

The grand total number of honeybees having taken a pesticide in an experimental colony is defined by the sum of the numbers of pre-existing (initial) adult bees and newborn ones for the duration of pesticide administration (CASE 1) or till extinction (CASE 2).

The following assumptions were made: (1) The age distribution of capped brood at an observation date is uniform between the first day when the cells of larvae are newly capped and the twelfth day when they eclose. (2) The number of adult bees that emerge from the pupae (capped brood) per day at a given day is one-twelfth of that of the capped brood at the last observation date before the day. (3) The total number of adult bees born between two successive observation dates is given by the product of one-twelfth of the number of capped brood at the former observation date and the number of days from the former to the latter observation date (4) The procedure in (3) is applied even when the number of days between two successive observation dates is greater than 12. (5) The number of capped brood at the time of the final pesticide administration or colony extinction is regarded as the number of adult bees having ingested the pesticide assuming that all the capped brood has already ingested the pesticide. (6) There is no newly capped brood after the middle of December because a queen stops laying eggs at the beginning of December in Japan

Now, we give an example of the procedure for estimating the grand total number of honeybees from Table 3 for the DF-Low/Pollen colony from July 9^th^ in 2011 to December 3^rd^ when the pesticide administration into the colony was stopped (CASE 1) under the above-mentioned assumptions. The number of initial adult bees is 2158; the total number of adult bees newborn between each two successive observation dates = (2556/12)(7-0) + (2377/12)(13-7) + (3207/12)(20-13), + (2743/12)(28-20) + (3480/12)(34-28) + (3217/12)(40-34) + (1933/12)(48-40) + (2099/12)(63-48) + (1343/12X70-63) + (1546/12)(77-70) + (2693/12)(82-77) + (2445/12)(90-82) + (2082/12)(104-90) + (1935/12)(113-104) + (2105/12)(118-113) + (2665/12)(132-118) + (1720/12)(140-132) + (1034/12)(148-140) = 27781; and the number of brood at the stop of the pesticide administration (December 3^rd^) is 840. That is, the grand total number of honeybees for the DF-Low/Pollen colony in CASE 1 till the stop of the pesticide (dinotefuran) administration on December 3^rd^ (during 148 days elapsed) is the sum (30779) of the initial bees (2158), the newborn ones (27781) and the final brood (840). Similarly, the grand total number of honeybees for the DF-Low/Pollen colony in CASE 2 is the sum (31076) of the initial bees (2158), the newborn ones (28918) obtained from the equation that 27781 + (840/12)(162-148) + 157 and the final brood (0).

CASE 1 is based on the assumption that the pesticide administered leads to the colony extinction while affecting honeybees (adult bees, brood etc.) only during the administration period of pesticide. CASE 2 is based on the assumption that it continues to affect all of honeybees in the colony till extinction even if it is discontinued halfway to be administered.

### Total intake of pesticide per bee

The total intake of pesticide per bee is defined by the division of the total intake of pesticide in a colony by the grand total number of honeybees having taken pesticide in the colony, which may be able to become an indicator to assess the collapse of a honeybee colony. Now, we give an example of the procedure for estimating the total intake of pesticide per bee for the DF-Low/Pollen colony in CASE 1: The total intake of the pesticide through pollen paste in a colony (DF-TIPP) is 1.8692 mg and the grand total number of honeybees (GTNH) is 30779 heads. Therefore, the total intake of pesticide per bee through pollen paste (DF-TIPP per bee) is 1.8692x10^6^/30779 = 60.73 ng/bee for CASE 1 of DF-Low/Pollen. Similarly, DF-TIPP per bee is 1.8692x10^6^/31076 = 60.15 ng/bee for CASE 2.

Table 5 shows the total intake of pesticide (dinotefuran) through sugar syrup in a colony (DF-TISS) or that through pollen paste (DF-TIPP) and the grand total number of honeybees in a colony (GTNH) obtained according to the procedure mentioned above, the total intakes of dinotefuran taken by individual bees during a period of the pesticide administration (CASE 1) or till extinction (CASE 2) through either sugar syrup or pollen paste in this work, and those taken through both sugar syrup and pollen paste in our previous work (Yamada *et al.*, 2012) for reference.

**Table 5.** Intake of foods (sugar syrup, pollen paste) and pesticide (dinotefuran)

|  | This Work:Experiment from 2011 to 2012 |  |  |  |  | Previous Work (Yamada et al., 2012):Experiment in 2010 |  |  |  |  |
| --- | --- | --- | --- | --- | --- | --- | --- | --- | --- | --- |
|  | Control | DF-Low/Syrup | DF-Low/Pollen | DF-High/Syrup | DF-High/Polle | Control | DF-Low | DF-Middle | DF-High | Control |
| Vehicle (food) to administer a pesticide |  | sugar syrup | pollen paste | sugar syrup | pollen paste |  | sugar syrup & pollen paste | sugar syrup & pollen paste | sugar syrup & pollen paste |  |
| Concentration in vehicle | 0 ppm of DF in both sugar syrup & pollen paste | 1 ppm of DF<sup>1)</sup> in sugar syrup | 0.565 ppm of DF in pollen paste<sup>2)</sup> | 10 ppm of DF in sugar syrup | 5.65 ppm of DF in pollen paste<sup>2)</sup> | 0 ppm of DF in both sugar syrup & pollen paste | 1 ppm of DF in sugar syrup & 0.333 ppm in pollen paste<sup>3)</sup> | 2 ppm of DF in sugar syrup & 0.667 ppm in pollen paste<sup>3)</sup> | 10 ppm of DF in sugar syrup & 3.33 ppm in pollen paste<sup>3)</sup> | 0 ppm of DF in both sugar syrup & pollen paste |
| Active Ingredient (AI) | pesticide-free | Dinotefuran | Dinotefuran | Dinotefuran | Dinotefuran | pesticide-free | Dinotefuran | Dinotefuran | Dinotefuran | pesticide-free |
| Pesticide administration (PA) | No | Continuous | Continuous | First one time | First one time | No | Continuous | Continuous | First one time | No |
|  |  | from Jul 9 to Oct 21 | from Jul 9 to Dec 3 | from Jul 9 to Jul 16 | from Jul 9 to Aug 6 |  | from Jul 18 to Oct 10 | from Jul 18 to Sep 17 | from Jul 18 to Jul 30 |  |
| Total intake of pesticide (dinotefuran) through sugar syrup in a colony (DF-TISS) or that through pollen paste (DF-TIPP) ; [mg] | 0.0000 | 4.208 from syrup | 1.8692 from pollen | 1.9 from syrup | 0.5257 from pollen | 0.0000 | 9.5 from syrup<br>0.47 from pollen | 9.98 from syrup<br>0.54 from pollen | 6.33 from syrup<br>0.5 from pollen | 0.0000 |
| **CASE 1 : Duration between the start of pesticide administration (PA) and the stop for an experimental colony or that of food feeding (FF) for a control colony** |  |  |  |  |  |  |  |  |  |  |
| Grand total number of honeybees in a colony (DTNH) during a period of PA for an experimental colony or that of FF for a control colony | 73074<br>from Jul 9 to Dec 17 in 2011 | 13543<br>from Jul 9 to Oct 21 in 2011 | 30779<br>from Jul 9 to Dec 3 in 2011 | 6544<br>from Jul 9 to Jul 16 in 2011 | 8078<br>from Jul 9 to Jul 16 in 2011 | 46260<br>from Jul 18 to Nov 21 in 2010 | 28500<br>from Jul 18 to Oct 10 in 2010 | 33931<br>from Jul 18 to Sep 17 in 2010 | 32277<br>from Jul 18 to Jul 30 in 2010 | 52212<br>from Jul 18 to Nov 21 in 2010 |
| Total intake of pesticide (dinotefuran) per bee during a period of PA for an experimental colony [AI ng/bee]<sup>5)</sup> ; DF-TISS/GTNH or DF-TIPP/GTNH | 0<br>from Jul 9 to Dec 17 in 2011 | 310.7 thru syrup<br>from Jul 9 to Oct 21 in 2011 | 60.73 thru pollen<br>from Jul 9 to Dec 3 in 2011 | 290.3 thru syrup<br>from Jul 9 to Jul 16 in 2011 | 65.08 thru pollen<br>from Jul 9 to Jul 16 in 2011 | 0<br>from Jul 18 to Nov 21 in 2010 | 333.3 thru syrup<br>16.49 thru pollen<br>from Jul 18 to Oct 10 in 2010 | 294.1 thru syrup<br>15.91 thru pollen<br>from Jul 18 to Sep 17 in 2010 | 196.1 thru syrup<br>15.49 thru paste<br>from Jul 18 to Jul 30 in 2010 | 0<br>from Jul 18 to Nov 21 in 2010 |
| Total intake of sugar syrup in a colony (TISS) during a period of PA for a experimental colony or that of FF for a control colony [g] | 26830<br>(pesticide-free) | 4208<br>(1 ppm of DF) | 18152<br>(pesticide-free) | 190<br>(10 ppm of DF) | 90<br>(pesticide-free) | 9500<br>(pesticide-free) | 9500<br>(1 ppm of DF) | 4987.5<br>(2 ppm of DF) | 633.33<br>(10 ppm of DF) | 9500<br>(pesticide-free) |
| Total intake of pollen paste in a colony (TIPP) during a period of PA for a experimental colony or that of FF for a control colony [g] | 4243<br>(pesticide-free) | 1459<br>(pesticide-free) | 3430<br>(0.565 ppm of DF) | 161<br>(pesticide-free) | 93<br>(5.65 ppm of DF) | 1500<br>(pesticide-free) | 1410<br>(0.3333 ppm of DF) | 810<br>(0.667 ppm of DF) | 150<br>(3.333 ppm of DF) | 1500<br>(pesticide-free) |
| TISS per bee during a period of PA for a experimental colony or that of FF for a control colony [g/bee] ; TISS/GTNH | 0.367 | 0.311 | 0.590 | 0.029 | 0.011 | 0.205 | 0.333 | 0.147 | 0.020 | 0.182 |
| TIPP per bee during a period of PA for a experimental colony or that of FF for a control colony [g/bee] ; TIPP/GTNH | 0.058 | 0.108 | 0.111 | 0.025 | 0.012 | 0.032 | 0.049 | 0.024 | 0.005 | 0.029 |
| **CASE 2 : Duration between the start of pesticide administration (PA) and the colony extinction (CE) for an experimental colony or that of food feeding (FF) for a control colony** |  |  |  |  |  |  |  |  |  |  |
| GTNH till CE for a experimental colony or during FF for a control colony<sup>4)</sup> | 73074<br>from Jul 9 to Dec 17 in 2011 | 13543<br>from Jul 9 to Oct 21 in 2011 | 31076<br>from Jul 9 in 2011 to Feb 16 in 2012 | 13457<br>from Jul 9 to Dec 17 in 2011 | 8582<br>from Jul 9 to Aug 6 in 2011 | 46260<br>from Jul 18 to Nov 21 in 2010 | 28500<br>from Jul 18 to Oct 10 in 2010 | 33931<br>from Jul 18 to Sep 17 in 2010 | 34642<br>from Jul 18 to Oct 30 in 2010 | 52212<br>from Jul 18 to Nov 21 in 2010 |
| Total intake of pesticide (dinotefuran) per bee till CE for an experimental colony [AI ng/bee]<sup>5)</sup> ; DF-TISS/GTNH or DF-TIPP/GTNH | 0<br>from Jul 9 to Dec 17 in 2011 | 310.7 thru syrup<br>from Jul 9 to Oct 21 in 2011 | 60.15 thru pollen<br>from Jul 9 in 2011 to Feb 16 in 2012 | 141.2 thru syrup<br>from Jul 9 to Dec 17 in 2011 | 61.26 thru pollen<br>from Jul 9 to Aug 6 in 2011 | 0<br>from Jul 18 to Nov 21 in 2010 | 333.3 from syrup<br>16.49 from pollen<br>from Jul 18 to Oct 10 in 2010 | 294.1 thru syrup<br>15.91 thru pollen<br>from Jul 18 to Sep 17 in 2010 | 182.7 thru syrup<br>14.43 thru paste<br>from Jul 18 to Oct 30 in 2010 | 0<br>from Jul 18 to Nov 21 in 2010 |
| Total intake of sugar syrup in a colony (TISS) during a period of PA for a experimental colony or that of FF for a control colony [g] | 26830<br>(pesticide-free) | 4208<br>(1 ppm of DF) | 18152<br>(pesticide-free) | 3660<br>(190g with DF of 10 ppm & the rest without pesticide) | 1130<br>(pesticide-free) | 9500<br>(pesticide-free) | 9500<br>(1 ppm of DF) | 4987.5<br>(2 ppm of DF) | 4750<br>(633.33g with DF of 10 ppm & the rest without pesticide) | 9500<br>(pesticide-free) |
| Total intake of pollen paste in a colony (TIPP) till CE for a experimental colony or during FF for a control colony [g] | 4243<br>(pesticide-free) | 1459<br>(pesticide-free) | 3430<br>(0.565 ppm of DF) | 1468<br>(pesticide-free) | 233<br>(93g with DF of 5.65 ppm & the rest without pesticide) | 1500<br>(pesticide-free) | 1410<br>(0.3333 ppm of DF) | 810<br>(0.6667 ppm of DF) | 1000<br>(150g of DF of 3.333 ppm & the rest without pesticide) | 1500<br>(pesticide-free) |
| TISS per bee till CE for a experimental colony or during FF for a control colony [g/bee] ; TISS/GTNH | 0.367 | 0.311 | 0.584 | 0.272 | 0.132 | 0.205 | 0.333 | 0.147 | 0.137 | 0.182 |
| TIPP per bee till CE for a experimental colony or during FF for a control colony [g/bee] ; TIPP/GTNH | 0.058 | 0.108 | 0.110 | 0.109 | 0.027 | 0.032 | 0.049 | 0.024 | 0.029 | 0.029 |

1) Dinotefuran as an active ingredient (A.I.)
2) Pollen paste was made of 15 parts of sugar syrup with dinotefuran and 10 parts of pollen without pesticide (dinotefuran) in weight: E.g., as pollen paste in DF-Low/Pollen is composed of 13 parts of sugar syrup with 1 ppm dinotefuran and 10 parts of pollen without dinotefuran, the concentration of dinotefuran in pollen paste becomes 1 [ppm]x13/23 ≈0.565 [ppm]
3) Pollen paste was made of 1 part of sugar syrup with pesticide (dinotefuran) and 2 parts of pollen without pesticide in weight: E.g., as pollen paste in DF-Low is composed of 1 part of sugar syrup with 1 ppm dinotefuran and 2 parts of pollen without dinotefuran, the concentration of dinotefuran in pollen paste becomes 1 [ppm]x1/3 ≈0.333 [ppm].
4) Estimated from the sum of the number of initial adult bees and that of newly-born adult bees by eclosion of capped brood.
5) Obtained by dividing the total intake of pesticide (dinotefuran) during feeding toxic food (DF-TISS or DF-TIPP) by the grand total number of honeybees (GTNH): E.g., for DF-Low/Pollen the total intake of pesticide (DF-TIPP) per bee from July 9 to December 3 is given by 1.8692 [mg] / 30779 ÷ 60.73 [ng]/bee in the case of CASE 1, and is similarly given by

The estimated GTNH in CASE 2 becomes more than that in CASE 1 because the pesticide is assumed to affect the colony for a longer period in CASE 2 including a pesticide-free period than in CASE 1 excluding the period. Consequently, the total intake of the pesticide (dinotefuran) per bee can be estimated lower in CASE 2 than in CASE 1 because the total intake of the pesticide for the duration of the experiment period in CASE 1 is same as that in CASE 2.

In both CASE 1 and CASE 2, the total intakes of dinotefuran per bee through sugar syrup (DF-TISS per bee) are roughly 300 ng/bee (DF-Low/Syrup, DF-High/Syrup) and those through pollen paste (DF-TIPP per bee) are about 60 ng/bee (DF-Low/Pollen, DF-High/Pollen) except for CASE 2 of DF-High/Syrup in this work.

In CASE 2 of DF-High/Syrup where the colony continued to survive for a long pesticide-free period before wintering after a short pesticide administration period, consequently the total intake per bee is about 140 ng/bee which is far below 300 ng/bee.

It can be seen from the results in CASE 1 of Table 5 in this work that DF-TISS per bee is 5.116 times (310.7/60.73) at low concentrations and 4.461 times (290.3/65.08) at high concentrations as much as DF-TIPP per bee, respectively. We can confirm that the intake of dinotefuran through pollen paste leading to the collapse of honeybee colony seems to be roughly one-fifth as much as that through sugar syrup in the case of CASE 1. Incidentally, the reason why DF-TIPP per bee in DF-High/Pollen (65.08 ng/bee) is slightly greater than that in DF-Low/Pollen (60.73 ng/bee) may be that the initial number of capped brood in DF-High/Pollen (6093) is more than double that in DF-Low/Pollen (2556) because brood take pollen paste (bee bread) in preference to sugar syrup (honey).

In our previous work conducted in 2010 (Yamada *et al.* 2012) where both sugar syrup and pollen paste with the pesticide were simultaneously administered to an experimental colony, the number of capped brood was given by the area ratio of the comb surfaces occupied with capped brood. We converted that into the number of capped brood from the average number (3249) of cells on every comb surface of all colonies. The average number was obtained as follows: The number of cells on each surface of every comb for all colonies was counted from photos without bees, which ranged from 3024 to 3432 and the mean value of each colony ranged from 3180 to 3318.

Figure 7 shows the total intakes of dinotefuran per bee for CASE 1 in this work and those in our previous work after converting DF-TIPP per bee into DF-TISS per bee with a 5.116-fold magnification at low concentrations, a 4.461-fold one at high concentrations and a 4.789-fold one at middle concentrations which is an average at low and high concentrations, respectively.

**Figure 7.**
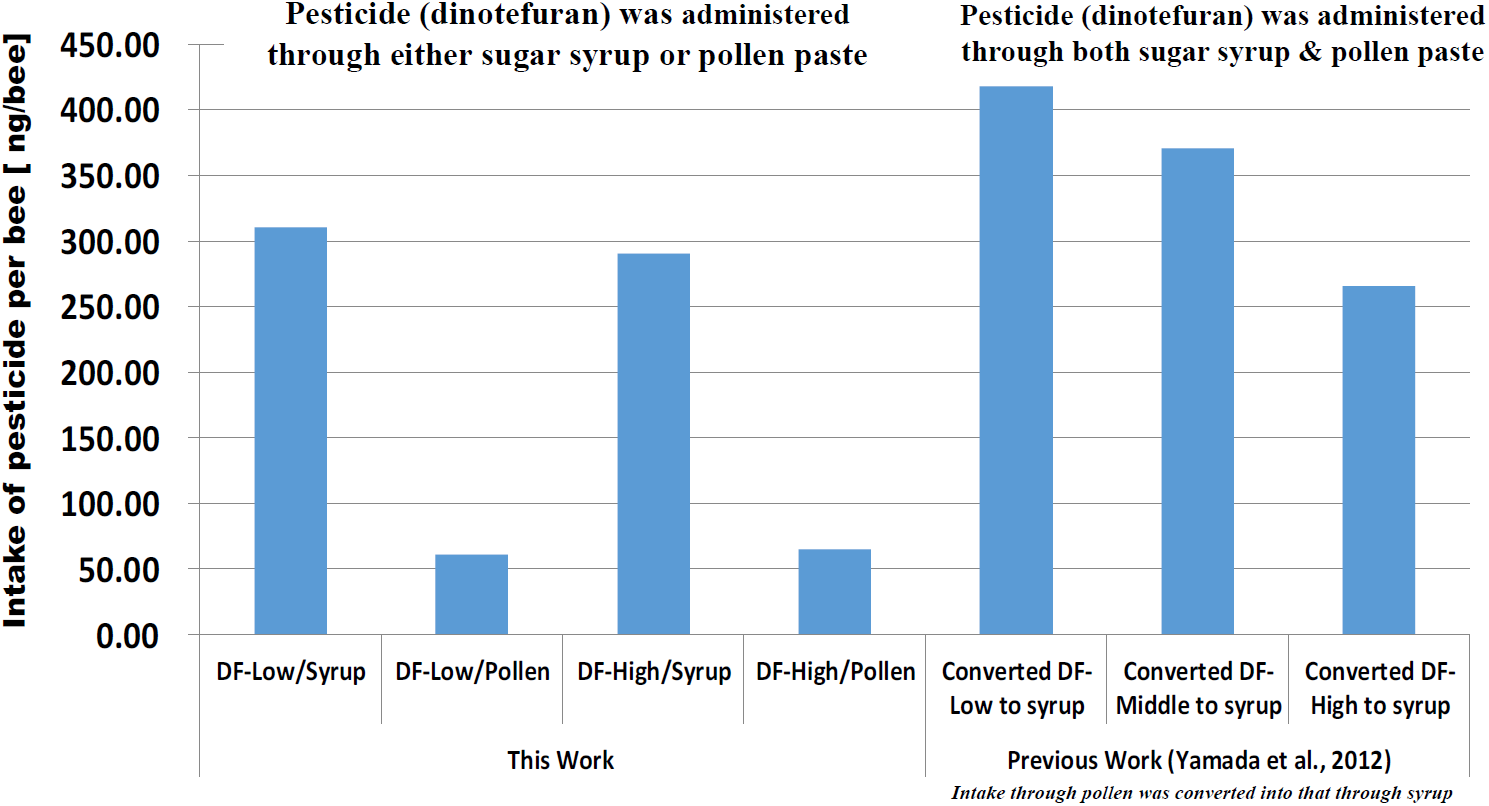
Total intake of dinotefuran per bee for CASE 1 The total intake of dinotefuran per bee in our previous work was obtained from the sum of dinotefura through sugar syrup after that through pollen paste which was converted into that through sugar syrup using the relation in this work between that through sugar syrup and that through pollen paste.

DF-TISS per bee in this work is in broad agreement with that after converting DF-TIPP per bee into DF-TISS per bee in previous work (Yamada *et al.* 2012). It is strange that DF-TISS per bee (ca. 300 ng/bee) is much more than LD_50_ ranging from 8 ng/bee to 61 ng/bee (US-EPA, 2004). Incidentally, the total intake through pollen paste (ca. 60 ng/bee) is on a comparable level with LD_50_.

### Total intake of each food per bee

Here, we turn our eyes to the total intake of each food (sugar syrup, pollen paste). The total intake of sugar syrup in a colony (TISS) or pollen paste in a colony (TIPP) in CASE 1 and that in CASE 2 are tabulated in Table 5. The total intake of each food per bee is obtained from dividing the TISS or TIPP by GTNH. TISS per bee and TIPP per bee for CASE 1 and CASE 2 in DF-Low/Syrup and DF-Low/Pollen-in this work and those in DF-Low in our previous work are almost equal to or more than those in each Control, respectively; that is, 0.311-0.59g/bee of sugar syrup, 0.108-0.111g/bee of pollen paste in DF-Low/Syrup, and 0.367 g/bee of sugar syrup and 0.058 g/bee of pollen paste in Control in this work; then, 0.333 g/bee of sugar syrup and 0.049 g/bee of pollen paste in DF-Low, and 0.182-0.205 g/bee of sugar syrup and 0.029-0.032 g/bee of pollen paste in Controls in our previous work.

On the other hand, those in CASE 1 in DF-High/Syrup and DF-High/Pollen in this work and those in DF-Middle and DF-High in our previous work are much less than those in each Control; that is, 0.011-0.029 g/bee of sugar syrup, 0.012-0.025g/bee of pollen paste in DF-High/Syrup and High/Pollen in this work; then, 0.020-0.147 g/bee of sugar syrup and 0.005 −0.024 g/bee of pollen paste to in DF-Middle and DF-High in our previous work.

In CASE 2 at high concentrations of dinotefuran, the total intakes of sugar syrup per bee in DF-High/Syrup (0.272 g/bee) and DF-High/Pollen (0.132 g/bee) in this work and that in DF-High (0.137 g/bee) in our previous work are much more than those in CASE 1, respectively; and then, those of pollen paste per bee in DF-High/Syrup (0.109 g/bee) and DF-High/Pollen (0.027 g/bee) in this work and that in DF-High (0.029 g/bee) in our previous work are much more than those in CASE 1, respectively. The reason for these increases is because the appetite of honeybees which have escaped death due to the administration of the pesticide (dinotefuran) came back during the feeding of pesticide-free foods. Though the improvement of their appetite during a pesticide-free period appears to be a proof of repellency at high concentrations, dinotefuran may only diminish their appetite rather than works repellently judging from enough intake of dinotefuran to lead to the colony extinction as compared with LD_50_.

## Discussion

### Effect of dinotefuran (neonicotinoid pesticide) on adult bees and brood

We will now discuss the effect of dinotefuran (neonocotinoid pesticide) on the number of adult bees and capped brood on the basis of experimental results from 2011, as shown in Tables 2 and 3 and Figures 1 to 3, proposing our plausible suppositions on the causal process where a honeybee colony leads to extinction.

High concentrations of dinotefuran, such as those in DF-High/Syrup and DF-High/Pollen, resulted in the presence of many dead bees near a hive and in the feeder within a few hours of administration. Considering that the concentration of pesticides sprayed on fields is about 10-fold higher than that in this work of the 2011 experiment, colonies can be presumed to collapse as follows: Foraging bees are instantly killed near the region where a high concentration of pesticide is sprayed because they take water, nectar, or pollen containing pesticide or contact with the pesticide. Instant death of foraging bees brings about a change in bee role, with house bees becoming foraging bees, thus resulting in the lack of house bees, and consequentially, in the collapse of the colony. The number of adult bees decreases markedly immediately after the temporary and brief administration of a high-concentration pesticide. Even after discontinuation of administration, their number continue to decrease, leaving a queen, small numbers of adult bees and brood and some amount of provisions with a very small number of dead bees in and around the hive. The reason why the number of adult bees decreased despite small number of dead bees is probably that foraging bees cannot return to the hive because of the nervous disorders or debility due to the toxicity of the pesticide ingested during their brood stage.

A queen did not die at a high concentration of the pesticide through sugar syrup (DF-High/Syrup) and died through pollen paste (DF-High/Pollen). The reason for this is broadly as follows: A queen is more susceptible to the pesticide through pollen paste than through sugar syrup because it can mainly take pollen. Even if a queen does not die by taking pesticides, it would lay only few eggs due to her reduced egg-laying capacity. Short time administration of pesticides with high concentrations leads to the instant death of many honeybees with acute toxic symptoms and even the subsequent pesticide-free administration finally leads to the colony extinction after presenting an appearance of a CCD because of the imbalanced colony structure, the reduced egg-laying performance of a queen and so on. Incidentally, it may be undeniable that toxic foods stored in a hive during pesticide administration continue to adversely affect honeybees even after the discontinuance of pesticide administration.

Continuous administration of a low-concentration pesticide (about 1% of the concentration recommended to exterminate stink bugs) caused colony extinction via an aspect of a CCD (DF-Low/Syrup) and a failure in wintering (DF-Low/Pollen). We found the following difference between the two vehicles: Colony collapse occurs earlier through sugar syrup than through pollen paste. This difference appears to be due to the total pesticide intakes which are 4.208 mg of dinotefuran in DF-Low/Syrup and 1.8692 mg in DF-Low/Pollen. The reason why a colony took less pesticide during a longer period through pollen paste than through sugar syrup is as follows: The intake period of pollen paste mainly taken by brood, whose developmental stage is shorter than that of adult bees, is shorter than the intake period of sugar syrup which is mainly consumed by adult bees. Add to this, the concentration of dinotefuran in pollen paste (0.565 ppm) is lower than that in sugar syrup (1 ppm). The reason why the colony failed in wintering in DF-Low/Pollen despite feeding pesticide-free foods before winter are probably because (1) the longevity of honeybees in DF-Low/Pollen could not increase due to the pesticide taken till then so much as that in Control could do with winter approaching and/or (2) honeybees continued to take toxic food stored in a hive.

### Comparison of the effect in toxicity of dinotefuran between this work and previous one

We now will consider the case where the insecticidal activity of dinotefuran through pollen paste is assumed to be higher than (about five times) that through sugar syrup. Figure 7 shows DF-TISS per bee for CASE 1 or DF-TIPP per bee in this work, and the sum of both the intake of dinotefuran through sugar syrup and that through pollen paste converted into that through sugar syrup in our previous work (Yamada et al., 2012), where both sugar syrup and pollen paste containing dinotefuran were administered into each experimental colony. When comparing the intakes through sugar syrup in this work with converted ones in our previous work, it can be seen from Figure 7 that they are on a comparable level with each other though the intakes in our previous work are somewhat higher than those in this work. These small differences may come from a state of colony such as the colony conditions and the environmental ones such as weather ones and seasonal ones, etc. We will consider whether the intake of food (sugar syrup, pollen paste) ingested by honeybees depends on the weather or not. As most of the total intake of dinotefuran is taken in a short period after the start of experiment as shown in Figure 6, it is enough to compare the weather during only a few weeks from the start of experiment between previous work and this one. Figure 8 shows the comparison between the weather during two weeks from the start of experiment in 2010 of our previous work (Yamada_*et al.*, 2012) and that in 2011 of this work in Noto District where we conducted the experimental in our apiary in Ishikawa Prefecture, Japan. As it can be seen from Figure 8 that there is little difference in temperature change between in 2010 and in 2011, the difference in the intake of dinotefuran seems not to come from the difference in weather (temperature) condition between two years.

**Figure 8.**
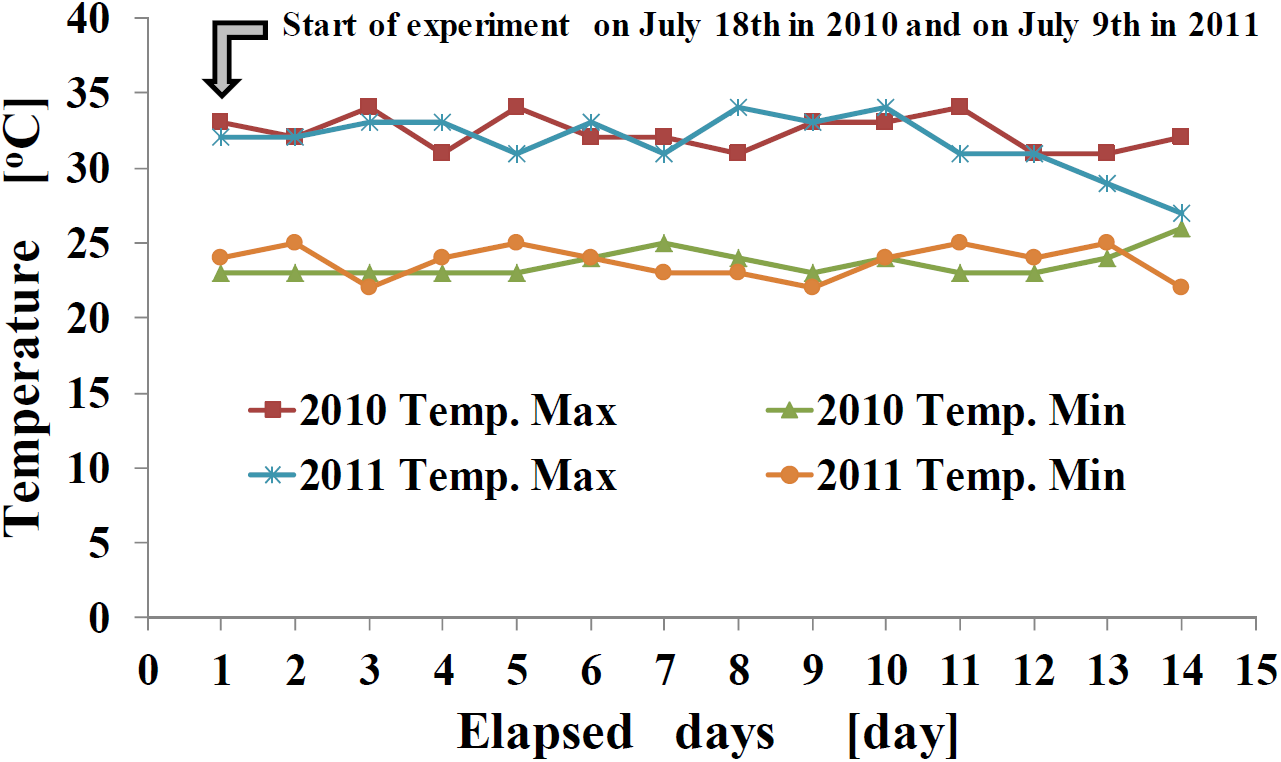
Change in temperature near the experimental site during two weeks from the start in 2010 (previous work) and in 2011 (this work)

We now will consider the case where the insecticidal activity of dinotefuran through pollen paste can be almost equal to that through sugar syrup under the assumption that a food which is stored in cells can be regarded as the apparent intake of a food. The longer a period of storage of a food is, the more the apparent intake of a food becomes. Bee bread (pollen paste) is usually stored in cells shorter than honey (sugar syrup). Supposing that pollen paste is immediately ingested by honeybees without storing it in cells, the intake of dinotefuran in our previous work can be given by the sum of both the intake through sugar syrup and that through pollen paste without conversion and then the sum in our previous work fairly approaches to the intake in this work (310.7 ng/bee in DF-Low/Syrup, 290.3 ng/bee in DF-High/Syrup). That is, the sum in our previous work for CASE 1 are 349.79 ng/bee in DF-Low, 310.01 ng/bee in DF-Middle and 211.59 ng/bee in DF-High. These values in our previous work become closer to those in this work.

We can confirm from the above two cases that the results on pesticide effect in our previous work have been substantially replicated through this work.

### Intake of food (sugar syrup, pollen paste) per bee

Before discussing the intakes of the pesticide (dinotefuran), we will begin by discussing the intakes of foods (sugar syrup, pollen paste) per bee in Table 5 which shows the details of experimental results for the two cases of CASE 1 and CASE 2. There are tabulated the total intake of pesticide through sugar syrup in a colony (DF-TISS) or that through pollen paste (DF-TIPP), the grand total number of honeybees in a colony (GTNH), the total intake of the pesticide (dinotefuran) per bee (DF-TISS per bee or DF-TIPP per bee), the total intake of sugar syrup in a colony (TISS), the total intake of pollen paste in a colony (TIPP), TISS per bee and TIPP per bee.

Compared with the estimated amount of sugar (about 1500 mg max. in total from larvae to foraging bees) reported by Rortais *et al.* (2005), which doubles when being converted into the amount of sugar syrup consisting of even amounts of sugar and water, every TISS per bee in Table 5 is lower than 3000 mg. TISS per bee ranges from 11 (DF-High/Pollen) to 590 mg/bee (DF-Low/Pollen) for CASE 1 and from 132 (DF-High/Pollen) to 584 mg/bee (DF-Low/Pollen) for CASE 2 in this work, and ranges from 20 (DF-High) to 333 mg/bee (DF-Low) for CASE 1 and from137 (DF-High) to 333 mg/bee (DF-Low) for CASE 2 in our previous work. Therefore, sugar syrup seems to be ingested after it is diluted more than five times with nectar gathered in fields.

TISS per bee in Control is 367 mg/bee in this work for both cases of CASE 1 and CASE 2. This value in Control is on a level with or less than TISSes in DF-Low/Syrup (311 mg/bee) and DF-Low/Pollen (590 mg) for CASE 1, and is also on a level with or less than TISSes in DF-Low/Syrup (311 mg/bee) and DF-Low/Pollen (584 mg/bee) for CASE2, and then is much higher than TISSes in DF-High/Syrup (29 mg/bee) and DF-High/Pollen (11 mg/bee) for CASE 1 and is higher than TISSes in DF-High/Syrup (272 mg/bee) and DF-High/Pollen (132 mg/bee) for CASE 2 in this work. TISSes per bee in Controls are 205 and 182 mg/bee in previous work for both cases of CASE 1 and CASE 2; these values in Controls are less than TISS in DF-Low (333 mg/bee) and are slightly higher than TISS in DF-Middle (147 mg/bee) for CASE 1 and CASE 2, and then are much higher than TISS in DF-High (20 mg/bee) for CASE 1 and are higher than TISS in DF-High (137 mg/bee) for CASE 2 in our previous work.

TIPP per bee in Control is 58 mg/bee in this work for both cases of CASE 1 and CASE 2. This value in Control is less than TIPPs in DF-Low/Syrup (108 mg/bee) and DF-Low/Pollen (111 mg) for CASE 1 and is also less than TIPPs in DF-Low/Syrup (108 mg/bee) and DF-Low/Pollen (110 mg/bee) for CASE2, and then is higher than TIPPs in DF-High/Syrup (25 mg/bee) and DF-High/Pollen (12 mg/bee) for CASE 1 and is less than TIPPs in DF-High/Syrup (109 mg/bee) and DF-High/Pollen (27 mg/bee) for CASE 2 in this work. TIPPs per bee in Controls are 32 and 29 mg/bee in previous work for both cases of CASE 1 and CASE 2. These values in Controls are less than TIPP in DF-Low (49 mg/bee) and are roughly on a level with TIPP in DF-Middle (24 mg/bee) for CASE 1 and CASE 2, and then are much higher than TIPP in DF-High (5 mg/bee) for CASE 1 and are on a level with TIPP in DF-High (29 mg/bee) for CASE 2 in our previous work.

It can be deduced that the neonicotinoid dinotefuran can be hardly repellent to honeybees at least at low concentrations where the intakes of foods are on a level with or more those in Control. Also at middle and high concentrations it seems probable that dinotefuran can hardly work repellently because honeybees continue to take toxic foods (sugar syrup, pollen paste) with dinotefuran enough for colony extinction. In spite of the presumption mentioned above, we cannot completely rule out a possibility that dinotefuran works repellently at middle or high concentrations, as it may appear to be due to a repellent effect that TISSes per bee and TIPPes per bee at middle or high concentrations are less than those at low concentrations.

Incidentally, each intake of foods in DF-High/Syrup and DF-High/Pollen for CASE 1 expectably becomes less than that for CASE 2, where pesticide-free foods were fed into an experimental colony after a short period of pesticide administration, respectively. Examining carefully the intakes of foods for CASE 2, the intakes of foods only for a pesticide-free period are 394 mg/bee of sugar syrup and 148 mg of pollen paste in DF-High/Syrup, and 459 mg/bee of sugar syrup and 62 mg of pollen paste in DF-High/Pollen, by estimating GTNH and the total intake of each food during a pesticide-free period. Though the intake of a toxic food is less in DF-High/Syrup or in DF-High/Pollen than the intake of food in Control, the intake of a nontoxic food increases up to the control level after the stop of pesticide administration.

Summering the above results, we can suggest the followings: (1) Dinotefuran seems hardly to work repellently for honeybees though it may cause anorexia at high concentrations: (2) At high concentrations of dinotefuran the provisioning of pesticide-free food can apparently recover the appetite but finally can allow leading to the extinction of colony after the exposure can cause a massive instant death and loss of appetite even for a short period of dosage: (3) The pesticide administered to a colony through sugar syrup seems to be diluted about five times (3000 mg /590 mg) in DF-Low/Pollen to about ten times (3000 mg/311 mg) in DF-Low/Syrup in this work. The dilution of toxic nectar will probably occurs actually in a hive in an apiary by adding pesticide-free nectar or pollen gathered in the fields. This means that high concentration neonicotinoid-pesticides in a field can act chronically after they are diluted with pesticide-free nectar, pollen, water, etc. which are foraged in the other fields and accumulated in a hive for a long period, because neonicotinoids is persistent (Yamada el al., 2012).

### Effect of pesticide intake through sugar syrup or pollen paste on a colony

We will now focus on the intake of pesticide per bee in Table 5. The discussion about the impact of dinotefuran (pesticide) on a honeybee colony is described below in two cases of CASE 1 and CASE 2 in comparison with our previous work (Yamada *et al.*, 2012).

Figure 7 shows TISS per bee is roughly about five times as TIPP per bee. If examined in detail, we can find the difference in the ratios of DF-TISS per bee to DF-TIPP per bee which are about 5.1-fold (310.7/60.73) at a low concentration (DF-Low/Syrup, DF-Low/Pollen), about 4.5-fold (290.3/65.08) at a high concentration (DF-High/Syrup, DF-High/Pollen) for CASE 1 in this work. Each average is about 300 ng/bee from (310.7+290.3)/2 through sugar syrup and about 63 ng/bee from (60.73+65.08)/2 and the ratio of the averages is about 4.8.

It has been proposed in the previous section of this paper that an about 4.8-fold difference between DF-TISS per bee and DF-TIPP per bee may come from the difference in efficacy or that in storage period. It is natural that this difference is attributed to the difference between the insecticidal activities through sugar syrup and those through pollen paste on a honeybee colony judging from the fact that pollen affects brood rather than adult bees. It can, however, be also considered that the difference comes from the difference between the storage period of sugar syrup and that of pollen paste stored in cells, judging from the fact that pollen paste is ingested for a short period (Gillian, 1979; DeGrandi-Hoffman *et al.*, 2013) though sugar syrup is ingested for a long period after diluted with nontoxic nectar gathered from fields and stored in a hive.

We now dare to try to explain the reason for the result that DF-TISS per bee which was much higher than the LD_50_ value of dinotefuran has not killed instantly all of honeybees in a colony, exploring the possibility to propose a new indicator for the assessment of persistent pesticide (dinotefuran) on colony failures such as a CCD and a wintering failure.

The reasons why DF-TISS per bee appears to be greater than the LD_50_ value of dinotefuran for an adult bee (8 ng/bee to 61 ng/bee), which is reported in Iwasa *et al.* (2004) and US-EPA (2004), seem to be as follows: (1) The difference between the LD_50_ and DF-TISS per bee comes from the difference between acute toxicity and chronic one which TISS possesses after diluted in cells with nontoxic nectar from fields. (2) A certain amount of dinotefuran in TISS is stored in combs as noxious honey, whose quantity changes with weather and seasonal conditions. (3) DF-TISS per bee is obtained from dividing DF-TISS by GTNH which is estimated with the error arisen from the assumptions described above. (4) The LD_50_ is determined from the quantity of a pesticide dosed out to an adult bee only once. In this work the intake of a pesticide (dinotefuran) in estimated from its quantity continuously taken by a honeybee through her life, without making an exception of a queen.

Here, we will discuss DF-TIPP per bee mainly for CASE 1. DF-TIPP per bee (60-65 ng/bee) in this work, where all of the experimental colonies finally have become extinct during experiments, appears to be slightly greater than and be approximately on a level with the LD_50_. We will examine the features on DF-TIPP and LD_50_ as follows: (1) Whereas honey is stored in cells for a long period, pollen is done only for a short period (Gillian, 1979; DeGrandi-Hoffman *et al.*, 2013). Therefore, pollen taken by honeybees seems to be hardly stored and then be directly and immediately ingested by them without little dilution. (2) The LD_50_ for an adult bee cannot be always applied for bood such a larva which takes mainly pollen. (3) DF-TIPP per bee is obtained from dividing DF-TIPP by GTNH which is estimated with the error arisen from the assumptions. (4) The LD_50_ is determined from the quantity of a pesticide dosed out to an adult bee only once. In this work the intake of a pesticide (dinotefuran) is estimated from its quantity continuously taken by a honeybee through her life, mainly during brood stages, without making an exception of a queen.

We now will consider whether TIPP in this work is valid for the usual amount of pollen taken by honeybees or not. TIPP per bee for CASE 1 in this work in Table 5 (as pollen paste, 58 mg in Control, 108 mg in DF-Low/Syrup, 111 mg in DF-Low/Pollen, 25 mg in DF-High/Syrup, 12 mg in DF-High/Pollen) seems to be reasonable because a honeybee consumes 112.5 mg to 195 mg of pollen in weigh in her life as reported by Crailsheim *et al.* (1992). Where 112.5 mg to 195 mg of pollen in weight is equivalent to 258.8 mg to 448.5 mg of pollen paste by multiplying the weight of pollen 2.3 times because pollen paste consists of one pollen and 1.3 sugar syrup. As the total consumption of pollen paste in her life is more than TIPP per bee in this work, most of TIPP in this work can be hardly stored and can be ingested till colony extinction because pollen is usually stored only for a short period (Gillian, 1979; DeGrandi-Hoffman *et al.*, 2013), while honeybees seem to supply a need by gathering pollen into a colony from fields.

Incidentally, the fact that DF-TISS per bee and DF-TIPP per bee are almost constant regardless of each concentration, respectively, suggests that dinotefuran may be probably poorly metabolized and mostly accumulated in the tissues of bees as pointed out in our previous work.

### Indicator for colony collapse

Although it is known that pesticides have an adverse effect on a honeybee in laboratory testing, the effect on a honeybee colony has been little elucidated in field testing. There are differences in susceptibility on a colony between field and laboratory tests to various factors such as environmental conditions, the behavior as a social insect, the storage of foods, the duration of the insecticidal effect and so on. LD_50_ is the amount of the substance (pesticide) required (usually per body weight) to acutely kill 50% of the test population during a given short time (24, 48 hrs, etc) where an individual compulsorily takes the substance (pesticide). Thus LD_50_ is an important indicator to evaluate the acute toxicity of a pesticide under limiting conditions but it can hardly assess the acute and chronic toxicities coexisting under practical field conditions, where the chronic toxicity is highly significant in persistent pesticides such as neonicotinoids. In field experiments conducted in a practical apiary the experimental concentration of a pesticide administered into a hive is changed by nectar, water and pollen with different concentrations gathered in fields. In a practical apiary honeybees are generally affected both acutely and chronically by pesticides. A honeybee colony collapses by various factors by which pesticides lead to the death of most honeybees due to acute toxicity, make honeybees feeble due to chronic toxicity to cause the infestations of mites and pathogens, cause them to lose their bearings due to chronic toxicity to cause them impossible to go back to their hive, cause the decrease in queen’s egg laying performance due to both acute toxicity and chronic one, cause the imbalance of populations among members such as foraging bees, house bees and brood by the death of particular honeybees due to both acute toxicity and chronic one and so on. We, however, cannot find an indicator to evaluate and assess the both acutely and chronically toxic effects of a pesticide on a honeybee colony, where exposure of honeybees to an acute toxicity causes an instant death or impede the return to their hive due to their physical weakening, and exposure to a chronic toxicity cause their disorientation because pesticides can affect the nerves. We will try to propose a new indicator (approach) to be able to assess the both effects of a persistent pesticide such as a neonicotinoid in a long-term field experiment.

Now, we will consider the possibility of DF-TISS per bee and DF-TIPP per bee as an indicator to assess the impact of persistent pesticide on a honeybee colony in a practical apiary. DF-TIPP per bee is almost the same and approximately 60 ng/bee both in CASE 1 and in CASE 2 regardless of concentration or period of pesticide administered into a honeybee colony. Surplus bee bread (pollen paste) is stored in cells and is consumed by honeybees in a relatively short span of time. This means that a honeybee colony seems to collapse when the intake of pesticide per bee is about 60 ng/bee regardless of acute or chronic toxicity if the pesticide is persistent and the efficacy is prolonged for a long period. It seems possible to use DF-TIPP per bee as an indicator to assess the colony collapse due to a persistent pesticide such as a neonicotinoid.

On the other hand DF-TISS per bee is much greater than DF-TIPP per bee and may be sometimes different between CASE 1 and CASE 2. The differences between CASE 1 and CASE 2 in DF-TISS per bee and between DF-TIPP per bee and DF-TISS per bee seems to be due to the storage of a pesticide such as a neonicotinoid which is hardly decomposed and is persistent. The amount of surplus honey (sugar syrup), which is able to be stored in cells for a long period, changes with the changes in environment such as the weather and the seasons. When DF-TISS per bee is greater than 60 ng/bee in a honeybee colony, the colony would be destined to collapse in due course of time even if it looks vigorous at a given time because honeybees in the colony continues to take stored toxic foods containing a persistent pesticide. If dinotefuran were easily decomposed and were not persistent, the toxicity in stored foods would lower with time, so that DF-TISS per bee of more than 60 ng /bee could not lead to the collapse in some cases.

Considering the wintering of a honeybee colony, honeybees, which have been newly born just before wintering, take toxic honey stored before winter in cells in a hive during wintering. If a honeybee continues to take only toxic honey containing dinotefuran during wintering till extinction under the assumptions that a honeybee takes the amount of sugar of 1500 mg in her life as reported by Rortais *et al.* (2005) and she dies by a dose of dinotefuran which is obtained from the estimation from DF-TIPP per bee in this work, we can assess the failure in wintering from the concentration of the pesticide included in honey in a hive. When thought is given to dinotefuran in this work, we can predict that a honeybee colony will collapse during wintering when honeybees continue to take toxic honey with a concentration higher than 40 ppb (60 [ng/bee] / 1500 [mg]) of dinotefuran stored before wintering as they live several times longer in winter than in other seasons and cannot take other foods besides stored honey in winter. It can be suggested from the above that a failure in wintering will be caused by stored honey with a low concentration of a pesticide. The concentration of a certain persistent pesticide in stored honey can probably constitute an indicator of failure in wintering once the total intake of the pesticide per bee through pollen paste till colony extinction is determined.

### Plausible story of neonicotinoids spayed in Japan

Now, we can deduce one of plausible scenarios in Japan when rice paddiy are sprayed with neonicotinoids which are systemic, persistent and high toxic, based on the facts obtained from our works.

After the rice seedlings treated by neonicotinoids are planted in a wet paddy in spring, the paddy is sprayed with neonicotinoids several times and the water in the wet paddy becomes contaminated with them. The contamination of the water lasts for a long period of time because of the persistent toxicity of them. When they are sprayed, the whole ecosystem in or near the rice paddy is contaminated by them. When their concentration is high, honeybees are killed instantaneously near the sprayed site. When they are low after diluted with the water in a rice paddy, honeybees are rarely killed on the sprayed site and carry the contaminated water with neonicotinoids into a hive, which have less repellent effect because they are odorless, tasteless and colorless. They use the water for cooling the hive by evaporation and for thinning honey to be fed to larvae. The contents of the hive such as honeybees, brood, eggs, honey, pollen, cells on combs, and so on are exposed to the toxicity of the contaminated water for a long period of time because neonicotinoids are persistent.

Similar situations to water arise in nectar and pollen. Honeybees import toxic nectar and toxic pollen into a hive, transfer them to the colony members and store them in cells after changing them into honey and bee bread, respectively. Honey can be stored in cells for a much longer period than pollen. Pollen is taken preferentially by brood and a queen and honey preferentially by adult bees. Toxicity in nectar and pollen is often diluted with nontoxic nectar imported from the other fields and stored in cells as mildly toxic honey and as mildly toxic bee bread made by kneading toxic pollen with nontoxic nectar (honey), respectively. Incidentally, nontoxic pollen is sometimes changed into toxic bee bread after kneaded with toxic nectar (honey) and nontoxic honey is also changed into toxic one when thinned with toxic water and fed to larvae.

Mildly toxic foods (honey, bee bread and water) are ingested by foraging bees, house bees, brood, and a queen, or stored in combs. When larvae that ingest mildly toxic bee bread, water and honey become foraging bees, they become unable to return to a hive because of either disorientation or exhaustion due to chronic toxicity. The egg-laying capacity of a queen declines through ingestion of mildly toxic foods, but she survives till a colony collapses. The death of many foraging bees creates an imbalance in the proportion of house bees, foraging bees, capped brood, and larvae in a colony, leading to colony collapse. Even if it does not collapse and appears vigorous, pesticides impede egg laying of a queen and result in a decrease in the bees’ immune strength, leading to infestations of mites, viruses, etc.

Put another way, when the pesticide concentration is high, the exposure to the pesticide causes an instantaneous death of bees due to acute toxicity. When the concentration is low, a CCD is caused by the exposure due to chronic toxicity. When the concentration is too low for a colony to become extinct before winter or honeybees take a low-concentration pesticide before winter, a failure in wintering is caused due to stored toxic foods or the aftereffect of the ingested pesticide even if toxic foods cannot be newly supplied into a colony during wintering. A CCD and a failure in wintering are phenomena characteristic of neonicotinoids, which are persistent, and can hardly be caused by organophosphates which are readily decomposed into their nontoxic components. We can apply this scenario on a rice paddy to an orchard and a farm.

## Conclusion

This study has reconfirmed the findings of our previous study (Yamada *et al.*, 2012) that a neonicotinoid such as dinotefuran causes the collapse and extinction of honeybee colonies after a honeybee colony has assumed the appearance of a CCD. The present results suggest that the insecticidal activity of dinotefuran in pollen paste (bee bread) on a honeybee colony is apparently roughly five times that in sugar syrup (honey or water), independently of pesticide concentration. On the other hand assuming that the difference between the intake of dinotefuran till the colony extinction through sugar syrup (honey) and that through pollen paste (bee bread) is due to the difference in a period of storage in a colony between honey and bee bread, the insecticidal activity of dinotefuran on a colony through sugar syrup can be considered to be almost equal to that through pollen paste. Further investigation on the difference in insecticidal activity on a colony is desired.

Although the intake of pesticide (dinotefuran) was enough to cause an instant death of a honeybee due to acute toxicity in this work judging from the LD_50_, we could find few dead bees near a hive and a colony dwindled away to nothing after assuming the appearance of a CCD due to chronic toxicity. The reason for a collapse of a colony due to chronic toxicity may be because sugar syrup and pollen paste containing the pesticide (dinotefuran) administered into a hive are diluted with nectar and pollen without pesticides in the experimental field which is controlled to be pesticide-free. Usually, foods for honeybees in certain fields (nectar, water, pollen) are contaminated by pesticides and ones in other fields are not done around an apiary. After the toxic foods are diluted with nontoxic ones and are stored in cells of a comb, they continue to adversely affect a colony as neonicotinoids are extremely persistent as compared with the other pesticides such as organophosphates. As a result, they cause a CCD and a failure in wintering due to chronic toxicity. Even when water and honey are temporarily contaminated by a neonicotinoid and they are stored in a hive for a long period of time, these phenomena may occur because of its persistency, but when pollen is temporarily contaminated by it, these phenomena may not always occur because of the short-term storage of bee bread (Gillian, 1979; DeGrandi-Hoffman *et al.*, 2013).

Pesticide intake per bee till the colony extinction is greater than the LD_50_ of dinotefuran in a honeybee, suggesting that the overall longevity of a colony, which behaves as one living creature, cannot be assessed by LD_50_, which is an indicator of acute toxicity to a honeybee. Pesticides impact honeybee colonies both acutely and chronically. In addition to LD_50_ expressing acute toxicity for a individual honeybee, an indicator providing information on honeybee colony strength is urgently needed in order to assess long-term pesticide-dosage effects on a complicated colony system exposed to toxicity ranging from chronic to acute.

This work suggests that the total intake of pesticide per bee through pollen paste could constitute an indicator of a persistent pesticide toxicity to cause a colony collapse and a failure in wintering in a practical apiary because dinotefuran may be probably poorly metabolized and mostly accumulated in the tissues of bees based on the result in this work that its insecticidal activity was independent from its concentration. By determining the intake through pollen paste to cause colony extinction and analyzing a concentration of the pesticide in honey before winter, we will probably judge the possibility of a failure in wintering because honeybees take mainly stored honey. Once we determine the total intake through pollen paste to cause colony extinction, we can judge the possibility of a CCD after measuring the concentrations of the pesticide in stored foods (honey and bee bread) and the average longevity of honeybees.

Based on this work and previous one (Yamada *et al.*, 2012), the following story will sound very convincing: When a pesticide is sprayed and dissolved in the water of a rice paddy or an orchard at low concentrations and is imported from fields to a colony by foraging bees, it continues to affect the colony for a long time and finally leads to a colony collapse or a failure of wintering. Even when a colony does not collapse and appears vigorous, insecticidal toxicity impedes the queen’s egg-laying capacity and reduces the bees’ immunity, shortening their lives or leading to mite infestations in the colony.

A neonicotinoid of very low concentration, which cannot be analytically detected, continues to gradually accumulate in the tissues of organisms over a long term and cause great harm to them. Thus, because of their high toxicity and relative nondegradability, neonicotinoids may be described as “agro-poisons” rather than agro-chemicals.

Since their creation in 1985, neonicotinoids have been commercially available from the early 1990s, and their advantages and disadvantages have been extensively discussed. Serious threats posed to nontarget animals, including human beings, have been revealed. For example, it is strongly suspected that neonicotinoids have caused a marked worldwide decline in freshwater arthropods, honeybees *(Apis mellifera)*, butterflies, red dragonflies, and sparrows and have exerted adverse effects on the human brain through their neurotoxicity (Beketov and Liess, 2008; JEPA, 2010; Kimura-Kuroda *et al.*, 2012a). Neonicotinoids may be poorly metabolized and mostly accumulated chronically in the tissues of bees at low concentrations. These may also represent a toxic threat to human beings (JEPA, 2010; Taira, 2012a, 2012b; Kimura-Kuroda *et al.*, 2012b), and we are fearful of a nightmare scenario in which Harm to honeybees can be applicable to human beings, and neonicotinoids may lead to the collapse of the Earth’s ecosystem.

## >Acknowledgments

We would like to express our gratitude to Ms. Rie Katsumata and Ms. Hiroko Nakamura, who made an enormous contribution to the accurate counting of the numbers of adult bees and capped brood by their arduous efforts. Here, we would also like to take this opportunity to thank all the individuals who assisted us in our research. This research was supported in part by Yamada Research Grant.

## Author Contributions

TY, KY and YY conceived and designed the experiments. TY and KY performed the experiments. TY and YY analyzed the data. TY and YY contributed reagents/materials/analysis tools. TY, YY and KY wrote the paper.

## References

Alaux, C Brunet, J-L; Dussaubat1, C; Mondet, F Tchamitchan, S Cousin, M Brillard, J Baldy, A Belzunces, L P; Conte, Y L (2010) Interactions between nosema microspores and a neonicotinoid weaken honeybees (Apis mellifera) Environmental Microbiology 12(3): 774–782 http://ONLINELIBRARY.WILEY.COM/doi:DOI/101111/J1462-2920.2009.02123.X/FULL

Beketov, M A; Liess, M (2008) Acute and delayed effects of the neonicotinoid insecticide thiacloprid on seven freshwater ARTHROPODS. Environmental Toxicology and Chemistry 27(2): 461–470. http://onlinelibrary.wiley.com/doi/1Q.1897/Q7-322R.1/abstract?deniedAccessCustomisedMessage=&userIsAuthenticated=false

Calderone, N W (2012) Insect Pollinated Crops, Insect pollinators and US agriculture: Trend analysis of aggregate data for the period 1992-2009. PLoS ONE 7(5): e37235. http://www.plosone.org/article/info%3Adoi%2F10.1371%2Fiournal.pone.0Q37235

Ciarlo, T J; Mullin, C A; Frazier, J L; Schmehl, D R (2012) Learning Impairment in Honey Bees Caused by Agricultural Spray Adjuvants. PLoS ONE 7(7): e40848 http://www.plosone.org/article/info%3Adoi%2F10.1371%2Fiournal.pone.004Q848

Colin, M E; Bonmatin, J M; Moineau, I Gaimon, C Brun, S Vermandere, J P (2004) A method to quantify and analyze the foraging activity of honeybees: Relevance to the sublethal effects induced by systemic insecticides. Archives of Environmental Contamination and Toxicology 47: 387–395 http://www.ncbi.nlm.nih.gov/pubmed/15386133

Cox-Foster, D L; Conlan, S Holmes, E C; Palacios, G Evans, J D; Moran, N A; Quan, P-L; Briese, T Hornig, M Geiser, D M; Martinson, V VAN Engelsdorp, D Kalkstein, A L; Drysdale, A Hui, J Zhai, J Cui, L Hutchison, S K; Simons, J F; Egholm, M Pettis, J S; Lipkin, W I (2007) A metagenomic survey of microbes in Honey Bee Colony Collapse Disorder. Science 318: 283–287 http://www.sciencemag.org/content/318/5848/283.short

Crailsheim, K Schneider, L H; Hrassnigg, N Buhlmann, G Brosch, U (1992) Pollen consumption and utilization in worker honeybees *(Apis Mellifera* Carnica): Dependence on individual age and function. Journal of Insect Physilogy 38(6): 409–419. http://www,sciencedirect,com/science/article/pii/00221910929Q117V

Degrandi-Hoffman, G Eckholm, B J; Huang, M H (2013) A comparison of bee bread made by Africanized and European bees (*Apis mellifera*) and its effects on hemolymph protein titers. Apidologie 44: 52–63. http://link,springer,com/article/1Q.10Q7%2Fs13592-Q12-Q154-9#page-1

El Hassani, A K; Dacher, M Gary, V Lambin, M Gauthier, M Armengaud, C (2008) Effects of sublethal doses of acetamiprid and thiamethoxam on the behavior of the honeybee (Apis mellifera). Archives of Environmental Contamination and Toxicology 54: 653661. http://link.springer.com/article/10.1007/s00244-007-9Q71-8#page-1

Gill, R J; RAMOS-Rodriguez, O Raine, N E (2012) Combined pesticide exposure severely affects individual-and colony-level traits in bees. Nature 491: 1Q5–1Q8. http://www.nature.com/nature/iournal/v491/n7422/abs/nature11585.html

Gillian, M (1979) Microbiology of pollen and bee bread: The yeasts. Apidologie 10(1): 43–53.

Henry, M Beguin, M Requier, F Rollin, O Odoux, J -F; Aupinel, P Aptel, J Tchamitchian, S Decourtye, A (2012) A common pesticide decreases foraging success and survival in honeybees. Science 336: 348–350. http://www.apidologie.org/articles/apido/pdf/1979/01/Apidologie004484351979101ART0006.pdf

Iwasa, T Motoyama, N Ambrose, J T; Roe, R M (2004) Mechanism for the differential toxicity of neonicotinoid insecticides in the honey bee. Crop Protection 23: 371–378. http://www.sciencedirect.com/science/article/pii/S0261219403002308

Japan Endocrine-Disruptor Preventive Action (JEPA) (2010) The threat of neonicotinoid pesticides on honeybees, ecosystems, and humans. http://kokumin-kaigi.sakura.ne.ip/kokumin/wp-content/uploads/Neonicotinoide.pdf THE JAPAN FOOD CHEMICAL RESEARCH FOUNDATION (JFCRF) (2014) Maximum Residue Limits (MRLs) of Agricultural Chemicals in Foods: Compositional Specification for Foods (Updated on January 30, 2014). The Japanese Positive List System for Agricultural Chemical Residues in Foods (Document released by Ministry of Health, Labour and Welfare). http://www.fFcr.or.jp/zaidan/FFCRHOME.nsf/pages/MRLs-p

Johnson, R M; Ellis, M D; Mullin, C A; Frazier, M (2010) Pesticides and honey bee toxicity-USA. Apidologie 41: 312–331. http://www.apidologie.org/articles/apido/pdf/2010/03/m09141.pdf

Kakuta, H Gen, M Kamimoto, Y Horikawa, Y (2011) Honeybee exposure to clothianidin: analysis of agrochemicals using surface enhanced Raman spectroscopy. Research Bulletin of Obihiro University 32: 31–36. http://iglobal.ist.go.ip/public/20090422/201102260879015920

Kimura-Kuroda, J Komuta, Y Kuroda, Y Hayashi, M Kawano, H (2012a) Nicotine-like effects of the neonicotinoid insecticides acetamiprid and imidacloprid on cerebellar neurons from neonatal rats. PLoS ONE 7(2): e32432. http://www.plosone.org/article/fetchObiect.action?uri=info%3Adoi%2F10.1371%2Fioumal.pone.0032432&representation=PDF

Kimura-Kuroda, J Komuta, Y Kawano, H (2012b) The effects of neonicotinoid pesticide on humans and mammals. Japanese Journal of Clinical Ecology 21(1): 46–56. http://www.asahikawa-med.ac.ip/dept/mc/healthv/isce/iice21146.pdf

Laurino, D Porporato, M Patetta, A Manino, A (2011) Toxicity of neonicotinoid insecticides to honey bees: laboratory tests. Bulletin of Insectology 64 (1): 107–113 http://smallbluemarble.org.uk/wp-content/uploads/2012/04/Laurino-2011.pdf

Lebuhn, G Droege, S Connor, E F; Gemmill-Herren, B Potts, S G; Minckley, R L; Griswold, T Jean, R Kula, E Roubik, D W; Cane, J Wright, K W; Frankie, G Parker, F (2013) Detecting Insect Pollinator Declines on Regional and Global Scales. Conservation Biology 27(1):113-120 http://onlinelibrary.wilev.com/doi/10.1111/i.l523-1739.2012.01962.x/abstract;isessionid=D2DC0A15D9DDF9099EA652C0318D6012.f04t01?deniedAccessCustomisedMessage=&userIsAuthenticated=false

Laycock, I Lenthall, K M; Barratt, A T; Cresswell, J E (2012) Effects of imidacloprid, a neonicotinoid pesticide, on reproduction in worker bumblebees (Bombus terrestris). Ecotoxicology 21: 1937–1945. http://link.springer.com/article/10.1007/s10646-012-0927-y#page-1

Lu, C Warchol, K M; Callahan, R A (2012) In situ replication of honeybee colony collapse disorder. Bulletin of Insectology 65(1): 99–106. http://www.bulletinoflnsectologv.org/pdfarticles/vol65-2012-099-106lu.pdf

Lu, C Warchol, K M; Callahan, R A (2014) Sub-lethal exposure to neonicotinoids impaired honey bees winterization before proceeding to colony collapse disorder. Bulletin of Insectology 67(1): 125–130. http://www.bulletinofinsectology.org/pdfarticles/vol67-2014-125-130lu.pdf

Marzaro, M Vivan, L Targa, A Mazzon, L Mori, N Greatti, M Toffolo, E P; Di BEmardo, A Giorio, C Marton, D Tapparo, A Girolami, V (2011) Lethal aerial powdering of honey bees with neonicotinoids from fragments of maize seed coat. Bulletin of Insectology 64(1): 119–126.http://www.moravbeedinosaurs.co.uk/neonicotinoid/aerial%20powdering%20of%20honev%20bees%20with%20neonicotinoids.pdf

Matsumoto, T (2013) Reduction in homing flights in the honey bee Apis mellifera after a sublethal dose of neonicotinoid insecticides. Bulletin of Insectology 66(1): 1–9. http://www.bulletinofinsectology.org/pdfarticles/vol66-2013-001-009matsumoto.pdf

Neumann, P Carreck, N L (2010) Honey bee colony losses. Journal of Apicultural Research 49(1): 1–6. http://www.ask-force.org/web/Bees/Neumann-Honev-Bee-Colonv-Losses-2010.pdf

Pettis, J S; Vanengelsdorp, D Johnson, J Dively, G (2012) Pesticide exposure in honey bees results in increased levels of the gut pathogen Nosema. Naturwissenschaften 99: 153–158 http://link.springer.com/article/10.1007/s00114-011-0881-1#page-1

Potts, S G; Biesmeijer, J C; Kremen, C Neumann, P Schweiger, O Kunin, W E (2010) Global pollinator declines: trends, impacts and drivers. Trends in Ecology and Evolution 25(6): 345–353 http://www.sciencedirect.com/science/article/pii/S0169534710000364

Rortais, A Arnold, G Halm, M -P; Touffet-BRIENS, F (2005) Modes of honeybees exposure to systemic insecticides: estimated amounts of contaminated pollen and nectar consumed by different categories of bees. Apidologie 36: 71–83. http://www.apidologie.org/articles/apido/pdf/2005/01/M4053.pdf

Schneider, C W; Tautz, J Grunewald, B Fuchs, S (2012) RFID Tracking of sublethal effects of two neonicotinoid insecticides on the foraging behavior of Apis mellifera. PLoS ONE 7(1): e30023. http://www.plosone.org/article/fetchObiect.action?uri=info%3Adoi%2F10.1371%2Fiournal.pone.0030023&representation=PDF?

Steinhauer, N A; Rennich, K Wilson, M E; Caron, D M; Lengerich, E J; Pettis, J S; Rose, R Skinner, J A; Tarpy, D R; Wilkes, J T; Vanengelsdorp, D (2014) A national survey of managed honey bee 2012-2013 annual colony losses in the USA: results from the Bee Informed Partnership. Journal of Apicultural Research 53(1): 1–18. http://dx.doi.org/10.3896/IBRA.1.53.1.01

Taira, K (2012a) Health effects of neonicotinoid insecticides. Part 1: Physicochemical Characteristics and Case Reports. Japanese Journal of Clinical Ecology 21(1): 24–34. http://www.asahikawa-med.ac.ip/dept/mc/healthy/isce/iice21124.pdf

Taira, K (2012b) Health effects of neonicotinoid insecticides. Part 2: Pharmacology, Application, Regulation, and Discussion. Japanese Journal of Clinical Ecology 21(1): 35–45. http://www.asahikawa-med.ac.jp/dept/mc/healthy/isce/iice21135.pdf#search=’%E3%83%8D%E3%82%AA%E3%83%8B%E3%82%B3%E3%83%81%E3%83%8E%E3%82%A4%E3%83%89%E7%B3%BB%E6%AE%BA%E8%99%AB%E5%89%A4%E3%81%AE%E3%83%92%E3%83%88%E3%81%B8%E3%81%AE%E5%BD%B1%E9%9F%BF+%E3%81%9D%E3%81%AE%EF%BC%92+%E5%B9%B3+%E8%87%A8%E5%BA%8A%E7%92%B0%E5%A2%83’

Taniguchi, T Kita, Y Matsumoto, T Kimura, K (2012) Honeybee colony losses during 2008-2010 caused by pesticide application in Japan. Journal of Apiculture 27(1): 15–27. http://www.dbpia.co.kr/Journal/ArticleDetail/2163286

Teeters, B S; Johnson, R M; Ellis, M D; Siegfried, B D (2012) Using video-tracking to assess sublethal effects of pesticides on honey bees (Apis melliferaL.). Environmental Toxicology and Chemistry 31(6): 1349–1354. http://onlinelibrary.wilev.com/doi/10.1002/etc.1830/full

United States Environmental Protection Agency (US-EPA) (2004) Pesticide Fact Sheet, Name of Chemical: Dinotefuran http://www.epa.gov/opp00001/chemsearch/regactions/registration/fsPC-04431201-Sep-04.pdf#search=’Pesticide+Fact+Sheet%2C+Name+of+Chemical%3A+Dinotefuran’

Van Der Sluijs, J P; Simon-Delso, N Goulson, D Maxim, L Bonmatin, J-M; Belzunces, L P (2013) Neonicotinoids, bee disorders and the sustainability of pollinator services. Current Opinion in Environmental Sustainability 5:1–13 http://dx.doi.org/10.1016/i.cosust.2013.05.007

Van Der Zee, R Brodschneider, R Brusbardis, V CHarriere, J-D; Chlebo, R Coffey, M F; Dahle, B Drazic, M M; Kauko, L Kretavicius, J Kristiansen, P Mutinelli, F Otten, C Peterson, M Raudmets, A Santrac, V Seppala, A Soroker, V Topolska, G VejsnÆs, F Alison Gray, A (2014) Results of international standardised beekeeper surveys of colony losses for winter 2012-2013: analysis of winter loss rates and mixed effects modelling of risk factors for winter loss. Journal of Apicultural Research 53(1): 19–34. http://dx.doi.org/10.3896/IBRA.1.53.1.02

Van Engelsdorp, D Caron, D Hayes, J Underwood, R Henson, M Rennich, K Spleen, A Andree, M Snyder, R Lee, K Roccasecca, K Wilson, M Wilkes, J; Lengerich, E Pettis, J (2012) A national survey of managed honeybee 2010-11 winter colony losses in the USA: Results from the Bee Informed Partnership. Journal of Apicultural Research 51(1): 115–124. http://www.ibra.org.uk/articles/US-honev-bee-winter-colonv-losses-2010-11

Van Tome, H V; Martins, G F; Lima, M A P; Campos, L A O; Guedes, R N C (2012) Imidacloprid-induced impairment of mushroom bodies and behavior of the native stingless bee melipona quadrifasciata anthidioides. PLoS ONE 7(6): e38406. http://www.plosone.org/article/fetchObiect.action7uriMnfo%3Adoi%2F10.1371%2Fiournal.pone.0038406&representation=PDF

Visser, A Blacquiere, T (2010) Survival rate of honeybee *(Apis mellifera)* workers after exposure to sublethal concentrations of imidacloprid. Proceedings of the Netherlands Entomological Society Meeting 21: 29–34. http://smallbluemarble.org.uk/wp-content/uploads/2012/04/029-034-VisserBlacquiere-2010.pdf

Whitehorn, P R; O’Connor, S Wackers, F L; Goulson, D (2012) Neonicotinoid pesticide reduces bumblebee colony growth and queen production. Science 336: 351–352. http://www.sciencemag.org/content/336/6079/351

Yamada, T Yamada, K Wada, N (2012) Influence of dinotefuran & clothianidin on a bee colony. Japanese Journal of Clinical Ecology 21(1): 10–23. http://dspace.lib.kanazawa-u.ac.ip/dspace/bitstream/2297/37606/1/SC-PR-YAMADA-T-10.pdf

Yang, E C; Chuang, Y C; Chen, Y L; Chang, L H (2008) Abnormal foraging behavior induced by sublethal dosage of imidacloprid in the honey bee (Hymenoptera: Apidae). Journal of Economic Entomology 101(6): 1743–1748. http://www.bioone.org/doi/abs/10.1603/0022-0493-10T6.1743

Yang, E.C; Chang, H C; Wu, W Y; Chen, Y W (2012) Impaired olfactory associative behavior of honeybee workers due to contamination of imidacloprid in the larval stage. PLoS ONE 7(11): e49472. http://www.plosone.org/article/fetchObiect.action7uriMnfo%3Adoi%2F10.1371%2Fiournal.pone.0049472&representation=PDF

